# De novo design of high-affinity miniprotein binders targeting *Francisella tularensis* virulence factor

**DOI:** 10.1101/2025.07.02.662053

**Authors:** Gizem Gokce-Alpkilic, Buwei Huang, Andi Liu, Lieselotte S.M. Kreuk, Yaxi Wang, Victor Adebomi, Yensi Flores Bueso, Asim K. Bera, Alex Kang, Stacey R. Gerben, Stephen Rettie, Dionne K. Vafeados, Nicole Roullier, Inna Goreshnik, Xinting Li, David Baker, Joshua J. Woodward, Joseph D. Mougous, Gaurav Bhardwaj

## Abstract

*Francisella tularensis* poses considerable public health risk due to its high infectivity and potential for bioterrorism. Francisella-like lipoprotein (Flpp3), a key virulence factor unique to Francisella, plays critical roles in infection and immune evasion, making it a promising target for therapeutic development. However, the lack of well-defined binding pockets and structural information on native interactions has hindered structure-guided ligand discovery against Flpp3. Here, we used a combination of physics-based and deep-learning methods to design high-affinity miniprotein binders targeting two distinct sites on Flpp3. We identified four binders for site I with binding affinities ranging between 24–110 nM. For the second site, an initial binder showed a dissociation constant (*K*_*D*_) of 81 nM, and subsequent site saturation mutagenesis yielded variants with sub-nanomolar affinities. Circular dichroism confirmed the topology of designed miniproteins. The X-ray crystal structure of Flpp3 in complex with a site I binder is nearly identical to the design model (Cα root-mean-square deviation: 0.9 Å). These designed miniproteins provide research tools to explore the roles of Flpp3 in tularemia and should enable the development of new therapeutic candidates.

## INTRODUCTION

*Francisella tularensis*, the causative agent of tularemia (commonly known as “rabbit fever”), is a highly infectious gram-negative intracellular pathogen. As few as 10–25 bacteria can initiate a severe infection after subcutaneous or aerosol delivery, leading to serious clinical outcomes such as pneumonia, sepsis, and multi-organ failure^1^. *F. tularensis* can leverage multiple immune evasion strategies to escape the phagosomes and replicate in the cytoplasm of macrophages and dendritic cells^2^. Due to its high risk for weaponization and significant threats from natural outbreaks and bioterrorism, the Centers for Disease Control and Prevention (CDC) classifies *F. tularensis* as a Tier 1 Select Agent and a Class A bioterrorism agent^3^. Despite considerable efforts, no approved vaccines are available for *F. tularensis*. While tularemia is currently treatable with antibiotics, such as fluoroquinolones, aminoglycosides, and tetracyclines, new medical countermeasures are needed to prepare for the natural and deliberate spread of emerging antibiotic-resistant strains of *F. tularensis*.

The outer membrane lipoprotein, Francisella-like lipoprotein (Flpp3), presents opportunities for developing novel research tools and therapeutic candidates against tularemia. It is a key virulence factor that plays critical roles in the spread and immune evasion of *F. tularensis*, and mutating the open reading frame encoding Flpp3 (FTT1416c) has been shown to significantly decrease its virulence^4^. Flpp3 plays a key role in establishing the initial bacterial infection and promoting bacterial progression by evading the immune system^5–8^. Flpp3 interferes with host immune responses, notably by playing roles in extending the lifespan of neutrophils through a TLR2-dependent mechanism that inhibits the apoptosis pathway, thereby promoting tissue destruction and bacterial dissemination^9^. Previous studies have shown that *F. tularensis* can bind and activate plasma plasminogen, contributing to extracellular matrix degradation and potential systemic dissemination. Several outer membrane lipoproteins, including Flpp3, have been identified as potential plasminogen-binding candidates, however, the specific role of Flpp3 in this process has not been established and remains to be experimentally confirmed^10^. While some Flpp3 functions are well studied, the full scope of Flpp3’s cellular localization, interactions with other proteins, and contributions to virulence remain unclear. Structurally, Flpp3 belongs to the bacterial lipoprotein family and shares low sequence and structural similarity to Bet v1 allergen proteins^11^. While it contains a single-helix membrane anchoring domain, studies suggest this may not be essential for membrane attachment, as lipidation can anchor Flpp3 to the outer membrane^12^. Flpp3 is also uniquely found in Francisella, making it an attractive target for therapeutic development, with the potential for developing ligands that disrupt its critical functions in immune evasion and bacterial dissemination.

The structure of Flpp3 has been characterized by X-ray Free-Electron Laser (XFEL) and Nuclear Magnetic Resonance (NMR) spectroscopy^8^, providing opportunities for the structure-guided design of Flpp3 inhibitors. However, there are still several challenges to targeting Flpp3 using rational approaches. First, no structurally-characterized binding partners of Flpp3 have been identified to date. Typically, the key interactions from such binding partners provide the basis for rationally designing new therapeutic candidates. Second, no deep pockets on the surface of Flpp3 are available for targeting with small molecules. While the NMR structure shows an internal cavity that could act as a small molecule binding pocket, this cavity was not observed in the XFEL structure^8^. Finally, the structural flexibility of Flpp3 also presents additional challenges for structure-guided design efforts. Despite these challenges, we reasoned that recent advances in physics-based and deep-learning (DL)-based design methods could enable the design of new Flpp3 binders. Such computational methods have recently been used to *de novo* design high-affinity miniprotein binders with diverse shapes and sizes against various therapeutic targets of interest^13–16^. Miniproteins, with their compact size, high structural stability, and capacity for high potency and specificity, offer a compelling choice for targeting membrane-bound proteins such as Flpp3.

Here, we leveraged *de novo* computational design methods to generate miniprotein binders targeting Flpp3. We identified multiple high-affinity binders with low nanomolar to picomolar binding affinities against two different sites of Flpp3, providing avenues to explore the cellular localization of Flpp3, its functional roles in infection, and to develop new therapeutic interventions against tularemia.

## RESULTS

### Computational Design of Flpp3-Binding Miniproteins

We used a combination of physics-based and DL-based methods to design, select, and optimize miniprotein binders against Flpp3. For initiating our design calculations, we used the structure of Flpp3 soluble domain (previously referred to as Flpp3sol^8^, PDB ID: 6PNY), which is composed of six β-sheets and two α-helices. Since the structural basis of Flpp3’s function as a virulence factor is largely unknown, particularly regarding the parts of the structure involved in interaction with the membrane or other interacting proteins to form potentially large molecular weight complexes, we decided to design minibinders against both faces of Flpp3 structure. We defined the electronegative face of Flpp3, dominated by the α-helices, and hypothesized to interact with the inner leaflet of the outer membrane as the site I (or ‘α site’). The opposite face formed by the β-sheets and exhibiting a relatively more electropositive character, was defined as the site II (or ‘β site’) (Figure 1A).

**Figure 1:**
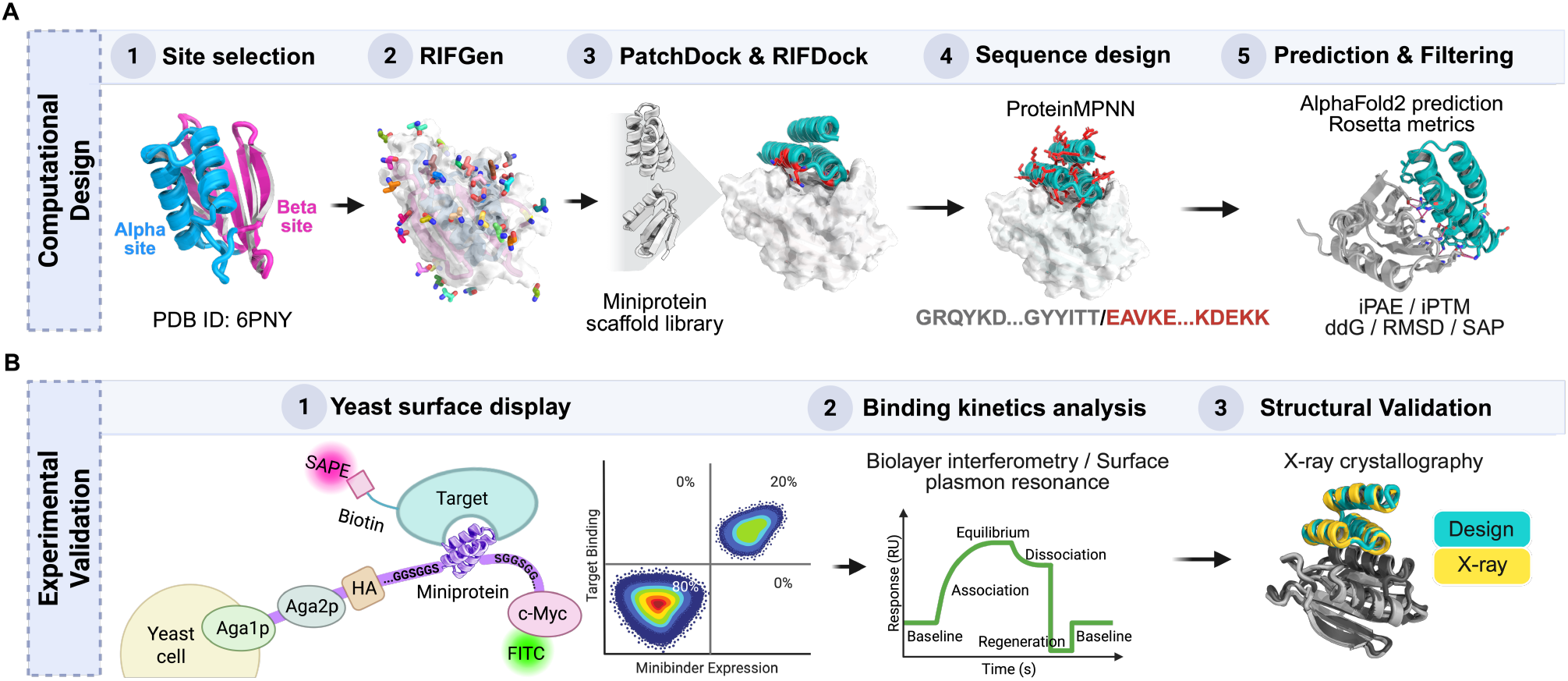
Design and screening pipeline for Flpp3-binding miniproteins. (A) Computational design workflow including site selection, RIFGen-based docking^43^, scaffold placement, ProteinMPNN-based sequence design, and structure prediction and filtering with AlphaFold2 and Rosetta. (B) Experimental validation pipeline including yeast surface display screening with FACS-based enrichment and next-generation sequencing of Flpp3-binding miniproteins, followed by binding kinetics measurement via biolayer interferometry or surface plasmon resonance, and structural / biophysical characterization.

To design miniprotein binders against the selected sites, we used a three-step computational pipeline (Figure 1). First, we used the Rosetta RIFDock^13^ approach to select miniprotein backbones with a high likelihood of making favorable interactions with Flpp3 from a pre-enumerated library of miniprotein scaffolds (Figure 1A). The initial step in the Rosetta RIFDock approach, RIFGen^13^, involves docking disembodied amino acids against the selected target site and calculating target binding energies. These docked amino acids then act as a rapid lookup table for estimating the target interaction energy achievable by a miniprotein scaffold based on its backbone coordinates and provide a means to select putative binders even before the sequence has been redesigned. We used a scaffold library of 43,724 pre-enumerated miniproteins composed of 25–65 amino acids and predicted to fold into diverse shapes. Next, we performed global shape complementarity docking using PatchDock^17^ to identify positions where the miniprotein scaffold and target achieve an optimal fit. The docking results were refined using RIFDock and a hierarchical search approach that uses positions generated by PatchDock as seeds for scaffold docking^17^. Overall, we generated and selected 500,000 docked conformations for further sequence design and *in silico* evaluation. Next, we used ProteinMPNN^18^, a deep learning-based tool for amino acid sequence design, to redesign the amino acid sequence of the selected minibinder backbones and improve the shape and chemical complementarity between the designed minibinders and Flpp3. Since ProteinMPNN does not allow backbone movement, we performed two iterative rounds of sequence design with ProteinMPNN followed by energy minimization using the Rosetta FastRelax protocol to allow for some backbone changes and achieve higher sequence diversity^19^. Finally, all designed minibinders were filtered based on the Rosetta metrics for interface quality and by re-predicting the structures of designed complexes using AlphaFold2 (AF2)^20^ to assess if designed models are confidently predicted to fold and bind as their designed conformations (Figure 1A). Newer structure prediction tools like AlphaFold3^21^ and RoseTTAFold-AA^22^ should provide a better and improved performance in filtering; however they were not available at the time and were not tried in this work. Specifically, we filtered designs based on AF2 pLDDT (>90 for α-site designs and >80 for β-site designs), interface PAE less than 6, Rosetta ddG values (<-40 kcal/mol for α-site designs and <-35 kcal/mol for β-site designs), and spatial aggregation propensity (SAP) scores (<30 for α-site designs and <35 for β-site designs). After the *in silico* filtering steps, we selected 15,000 α-site design models and 8,817 β-site design models for experimental screening and characterization.

### Yeast Display Screening and Biochemical Characterization

We screened the selected α-site and β-site miniproteins for binding to Flpp3 using yeast surface display (YSD) and fluorescence-activated cell sorting (FACS). Synthetic oligonucleotides encoding for the designed minibinders were commercially purchased and cloned into a pETCON3 expression vector for display on the surface of the yeast cells (see Methods for details) (Figure 1B). The α-site and β-site design libraries were ordered, cloned, and screened separately. After an initial expression sort to collect the yeast cells expressing the designed minibinders, a round of binding sort was conducted with 1 µM of Flpp3 to collect yeast cells that bind fluorescently labeled streptavidin-tetramerized Flpp3. We observed a clear binding signal at 1 µM for both α-site and β-site libraries (Supplementary Figure 1). To identify and select for the higher-affinity binders among this initial binding population of cells, the binding cells from the initial binding sorts were grown and sorted again with decreasing concentrations of monomeric Flpp3: 1000 nM, 100 nM, 10 nM, and 1 nM for the α-site library and 1000 nM, 300 nM, 100 nM, 30 nM for the β-site library (Figure 2A). The α-site library showed a clear population of Flpp3-binding cells even at 1 nM, but the signal for the β-site library screened with 30 nM Flpp3 returned to levels comparable to the negative control (*i*.*e*., sorting with no Flpp3) (Figure 2B). After sequencing the binding cells from different sorting experiments using next-generation sequencing (NGS) and calculating the enrichment levels of each sequence across the experiments, we identified four promising binder candidates for the α-site (ASD1–4) and one additional binder (BSD1) for the β-site.

**Figure 2.**
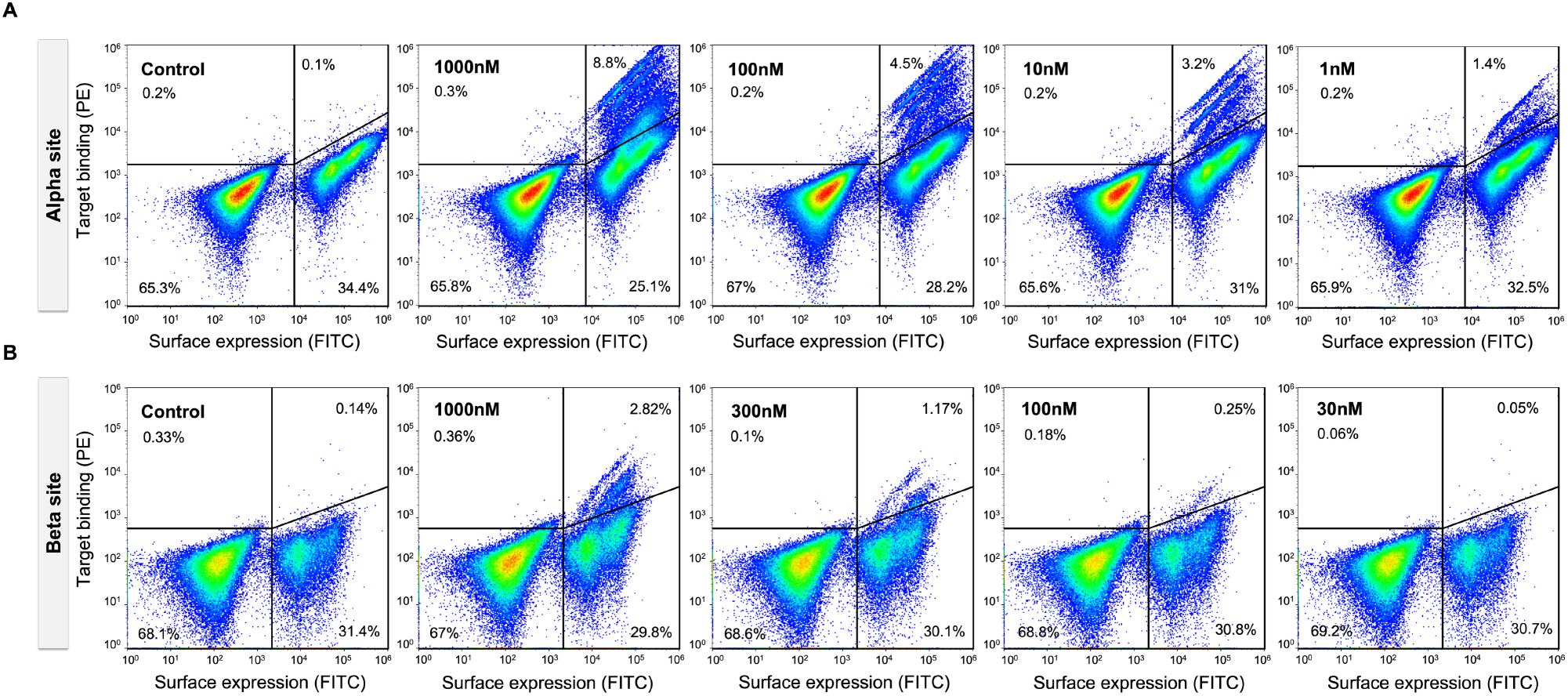
Yeast surface display screening of α-site and β-site libraries against Flpp3. (A) Binding profiles of the α-site library at decreasing concentrations of Flpp3 (1000 nM, 100 nM, 10 nM, and 1 nM). A distinct population of Flpp3-binding cells is observed even at the lowest concentration (1 nM), indicating high-affinity binders. (B) Binding profiles of the β-site library at decreasing concentrations of Flpp3 (1000 nM, 300 nM, 100 nM, and 30 nM). The binding signal at 30 nM is comparable to the negative control (no Flpp3), suggesting weaker binding relative to the α-site.

The design models for all five binders from YSD screening have a three-helix topology with two helices making extensive contacts with the Flpp3 surface (Figure 3A). Each putative binder demonstrated excellent computational metrics, such as AF2 iPAE < 3.5, and Rosetta ddG < −40 kcal/mol (Figure 3A), indicating favorable binding interactions. To confirm the binding in solution (in addition to the binding on yeast surface) and determine binding affinities, we expressed the five putative binders in *E. coli* and purified them using immobilized metal affinity chromatography (IMAC) and size-exclusion chromatography (SEC). All binders expressed well with a single peak corresponding to the monomer size on SEC (Figure 3B). Next, we used Biolayer interferometry (BLI) to confirm the binding of the miniproteins to the biotinylated Flpp3. Biotinylated-Flpp3 was immobilized at 50-100 nM concentrations on streptavidin-coated biosensors, and binding kinetics were measured by monitoring association and dissociation across a series of miniprotein concentrations in BLI assay buffer. All four α-site designs showed binding to Flpp3 with binding affinities (*K*_*D*_) ranging between 24–110 nM (Figure 3C), with designs ASD1, ASD2, and ASD3 showing *K*_*D*_ < 50 nM and the design ASD1 showing the best binding affinity (*K*_*D*_: 24 nM) for the α-site. The β-site design, BSD1, also showed promising binding to Flpp3 with a *K*_*D*_ of 190 nM.

**Figure 3:**
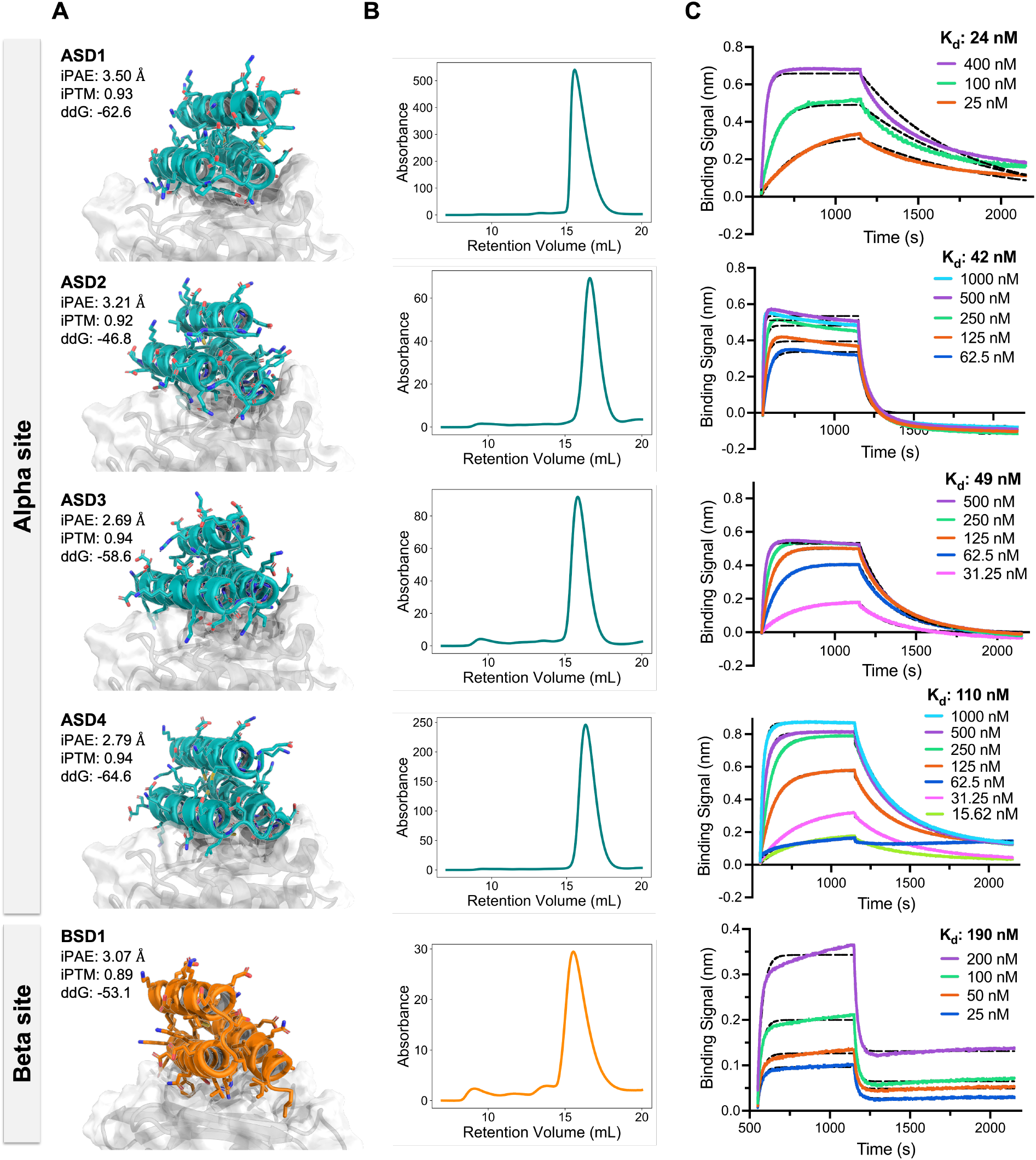
Design models and biochemical characterization of Flpp3-binding miniproteins. (A) Design models of the five binders (four for α-site in teal and one for β-site in orange). Flpp3 shown in gray. (B) Size-exclusion chromatography (SEC) profiles of the purified binders expressed in *E. coli*, demonstrating monodisperse peaks consistent with the monomer size of the minibinders. (C) Bio-layer interferometry (BLI) binding curves for the designed binders against biotinylated Flpp3. Calculated K_D_ for each binder reported on the plots.

Computational models for all four α-site binders (ASD1-4) have similar structural features and show similarity in binding mode and interactions. ASD1 and ASD4 show very similar three-helix fold, binding mode, and interface contacts. ASD2 also has the same three-helix fold and targets the same region of Flpp3, but shows a slightly shifted helix orientation (Figure S1). Despite differences in their overall amino acid sequences, ASD1, ASD2, and ASD4 are predicted to engage a similar region on Flpp3 with subtle differences in the types of interface contacts. ASD1 and ASD4 both include an Asp residue that forms a polar interaction with Tyr42, Phe43, and Arg102 of Flpp3, while ASD2 places a Trp at the same position. ASD2 and ASD4 also share an Arg and a Glu at an adjacent site, while ASD1 features a Tyr at the same location. ASD3 adopts a shifted backbone orientation and uses a different set of interface residues. Sequence comparisons show moderate similarities among ASD1, ASD2, and ASD4 (25–40% pairwise sequence identity), while ASD3 is more divergent from other designs (20–25% pairwise identity) (Figure S2). The design model for the β-site binder BSD1 shows an extensive network of hydrophobic and polar interactions. Together, these results highlight the ability of computational design to generate high-affinity binders against multiple sites of a selected target.

### Optimization of the β-site Binders

Since the β-site binders generally demonstrated lower binding signal and affinities than α-site binders during YSD and BLI experiments, we next set out to improve the affinity of the β-site binder BSD1 using site saturation mutagenesis (SSM). A pooled library of 1,046 BSD1 variants was generated by mutating each residue of BSD1 to 19 other canonical amino acids. Oligonucleotides encoding this variant library were commercially purchased, cloned into yeast cells, and screened for binding to Flpp3 using FACS as described earlier. In contrast to the original library of β-site designs (Figure 2B), which did not show a binding signal at 30 nM of Flpp3, the SSM-optimized library showed a clear binding signal even during the sorting runs with 1 nM of monomeric Flpp3, indicating the presence of improved binders (Figure 4A).

**Figure 4:**
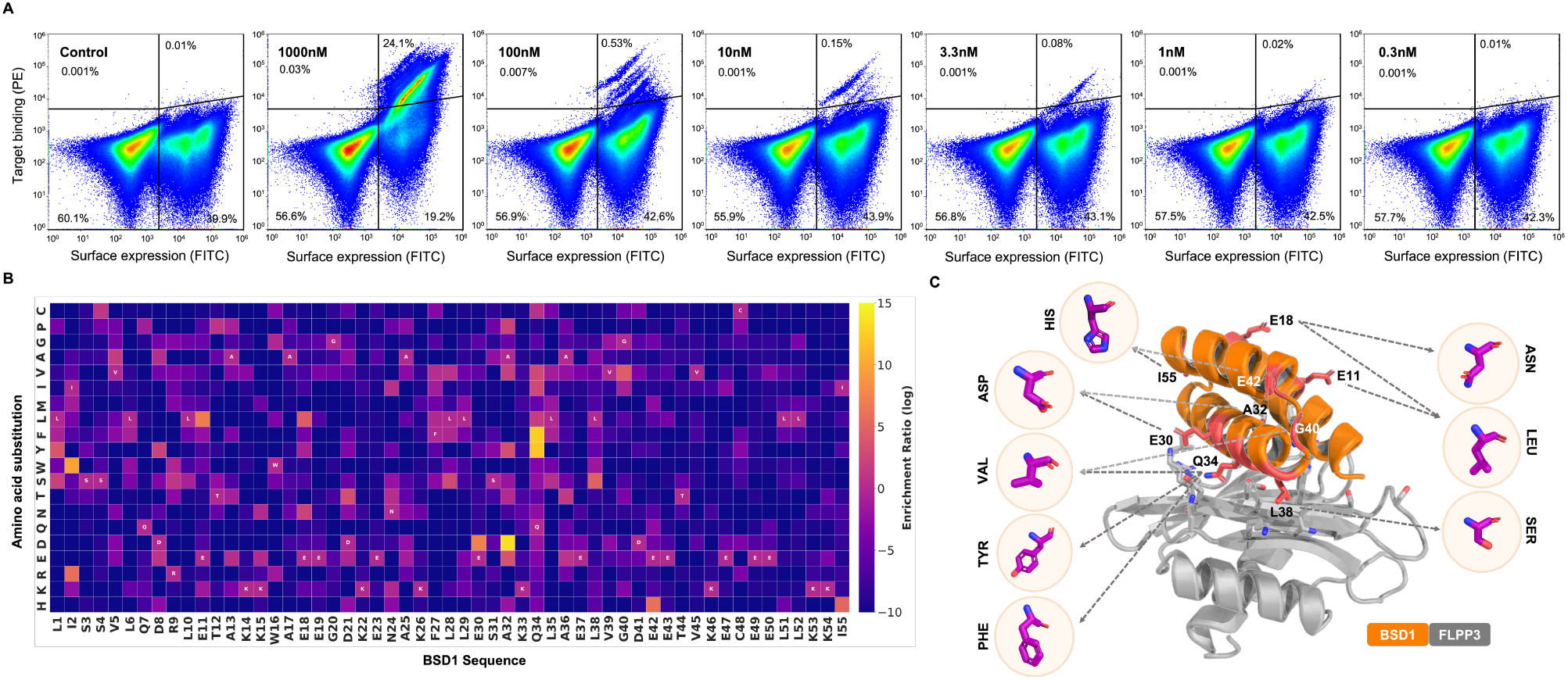
Site saturation mutagenesis optimization of β-site binder BSD1. (A) Fluorescence-activated cell sorting plots show binding of BSD1 variants at decreasing concentrations of monomeric Flpp3 (1000 nM to 0.3 nM). (B) Mutational enrichment heatmap of BSD1 SSM variants. The heatmap represents log-transformed enrichment ratios of amino acid substitutions at each position of BSD1, derived from next-generation sequencing (NGS) data across multiple FACS-based binding sorts. Enrichment values were calculated by normalizing amino acid frequencies within the selected population relative to the naive library. Positions with high enrichment (yellow) correspond to beneficial mutations that enhance Flpp3 binding, whereas positions with low enrichment (purple) indicate substitutions that reduce binding. (C) Structural mapping of enriched substitutions onto the BSD1. Selected residues on BSD1 are shown with side chains highlighted in coral, representing sites where substitutions were most strongly enriched in the YSD screen. Magenta callouts show example side chains predicted to improve binding affinity, based on enrichment data. These mutations primarily localize to the binder’s interface with Flpp3, suggesting their role in stabilizing key interactions.

**Figure 5:**
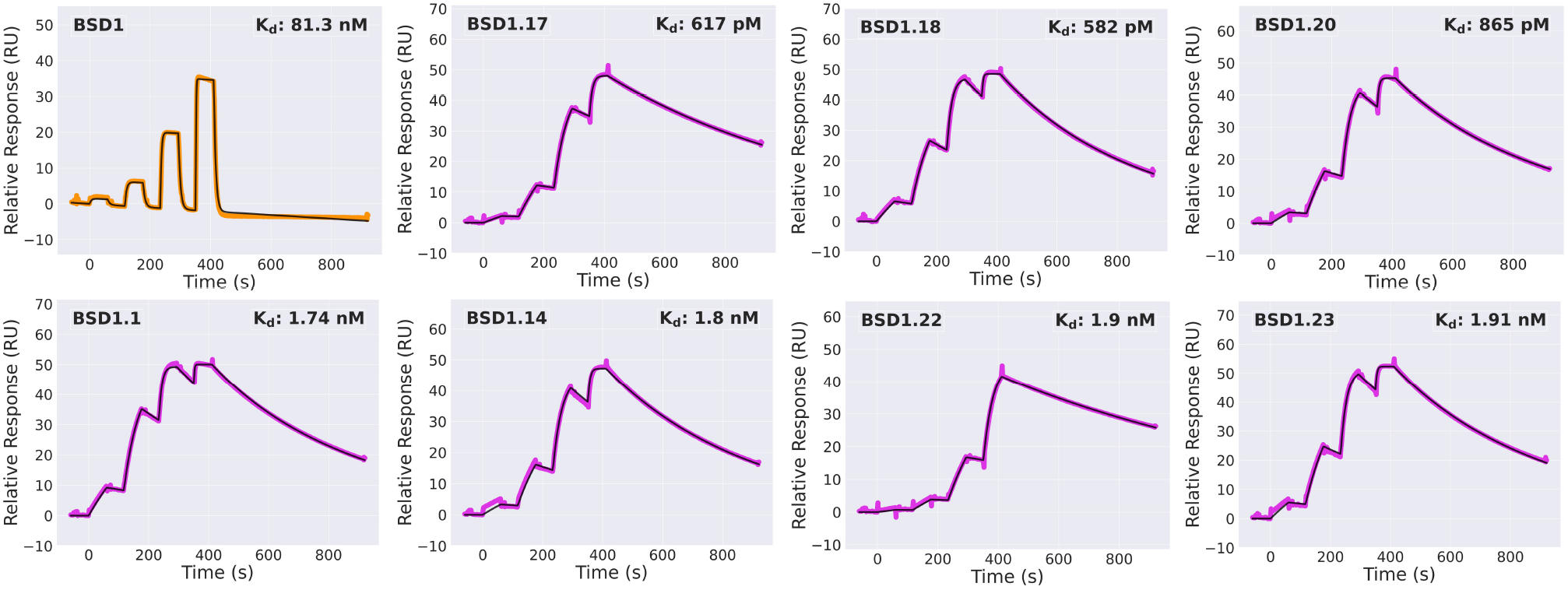
Determination of binding affinities for BSD1 and its selected variants using SPR. SPR sensorgrams from a 4-point single cycle kinetics experiment (5-fold dilution, highest concentration ranging from 0.05 µM to 0.43 µM depending on the minibinder). Experimental data are shown in orange/magenta, and global fits are shown with black lines.

Sequencing data from the SSM library sorting supports the predicted binding mode of BSD1, with high conservation across positions making substantial interface contacts in the design model (Figure 4C). Additionally, the SSM data also suggested nine amino acid positions where mutations could potentially improve the binding affinity. Based on this data, we selected twelve single-mutation variants, seven double-mutation variants, and five triple-mutation variants of BSD1 for further characterization (Table 1). All variants (BSD1.1–BSD1.24) were expressed in *E. coli*, purified using IMAC and SEC, and assessed for Flpp3 binding using surface plasmon resonance (SPR). One variant (BSD1.21) did not express well and was not pursued further. The native design showed a K_D_ of 81 nM in SPR single-cycle kinetics experiments (compared to 190 nM by BLI). 18 out of the 22 tested variants showed better *K*_*D*_ than the original binder BSD1, with 13 variants yielding *K*_*D*_ values less than 10 nM. Notably, single-residue mutation of an interface-facing Gln34 to Phe in BSD1.1 improved the binding affinity from 81 nM to 1.7 nM. Two double mutation variants (Q34F/I55H, and Q34F/L38S) and one triple-mutation variant (Q34F/E11L/E30D) showed sub-nanomolar binding affinities, with the two best variants, BSD1.17 and BSD1.18, displaying *K*_*D*_ of 582 pM and 617 pM, respectively. Both BSD1.17 and BSD1.18 are still predicted to fold and bind similarly to BSD1 by AlphaFold3 (Figure S8). Notably, all variants with sub-nanomolar binding affinity include the Q34F mutation, highlighting the importance of this particular interface mutation towards achieving improved binding affinity.

**Table 1.** Binding affinities of BSD1 variants against Flpp3. Summary of single and combinatorial amino acid substitutions introduced into the parental minibinder BSD1. Binding affinities (in nM) were measured by SPR. Variants are sorted by name, and substitutions are listed relative to BSD1.

| *Name* | *Substitution(s)* | *Binding affinity (nM)* |
| --- | --- | --- |
| BSD1 | Original binder | 81.30 |
| BSD1.1 | Q34F | 1.74 |
| BSD1.2 | Q34Y | 5.72 |
| BSD1.3 | Q34V | 12.84 |
| BSD1.4 | E11L | 9.50 |
| BSD1.5 | E18N | 249.80 |
| BSD1.6 | E18L | 8.72 |
| BSD1.7 | E30D | 17.62 |
| BSD1.8 | A32D | 8.24 |
| BSD1.9 | L38S | 367.60 |
| BSD1.10 | G40V | 59.40 |
| BSD1.11 | E42H | 87.35 |
| BSD1.12 | I55H | 31.46 |
| BSD1.13 | Q34F, E11L | 1.67 |
| BSD1.14 | Q34F, E30D | 1.80 |
| BSD1.15 | Q34F, A32D | 125.10 |
| BSD1.16 | Q34F, E42H | 2.73 |
| BSD1.17 | Q34F, L38S | 0.62 |
| BSD1.18 | Q34F, I55H | 0.58 |
| BSD1.19 | E11L, E30D | 13.77 |
| BSD1.20 | Q34F, E11L, E30D | 0.86 |
| BSD1.21 | Q34F, E11L, E42H | N/A |
| BSD1.22 | Q34F, E11L, L38S | 1.90 |
| BSD1.23 | Q34F, E30D, I55H | 1.91 |
| BSD1.24 | E11L, E30D, E42H | N/A |

### Thermal Stability and Secondary Structure Characterization by Circular Dichroism

Next, we used circular dichroism (CD) to assess the thermal stability and obtain low-resolution structural information for the designed minibinders. We measured the CD spectra for the two best α-site designs (ASD1 and ASD2), the best β-site design from the first round (BSD1), and the best variant of BSD1 after SSM maturation (BSD1.18). Both α-site designs, ASD1 and ASD2, show a spectrum consistent with an all-helical topology at 25°C (Figure S7A, left panel). The thermal melt curves show that ASD1 and ASD2 unfold when heated to 95°C, with melting temperatures of 55°C and 65°C, respectively (Figure S7A, right panel). However, both ASD1 and ASD2 refold to the original fold upon cooling to 25°C (Figure S7A, left panel). Similar to α-site binders, the β-site binder BSD1 also shows spectra typical for an all-helical topology at 25°C, suggesting that it adopts the designed topology (Figure S7B, left panel). The minimal CD change observed for both β-site binders during thermal melting experiments indicates that they remain folded across the tested temperature range (25-95°C), suggesting a hyperstable structure with a melting temperature beyond 95°C. These data suggest that both BSD1 and the SSM-optimized β-site binder BSD1.18 are more thermostable than the α-site binders. Overall, the α-site and β-site binders adopt the designed topology and refold to the designed fold even after thermal denaturation up to 95°C and subsequent cooling.

### Structure Determination of Flpp3-ASD1 Complex by X-ray Crystallography

To confirm the three-dimensional structure and binding mode of our designed minibinders, we pursued crystallization trials for several α-site and β-site designs in complex with Flpp3. Among these, only the highest-affinity α-site binder, ASD1, yielded crystals suitable for X-ray diffraction. We determined the X-ray crystal structure of the ASD1–Flpp3 complex at 2.37 Å resolution. The structure matches very closely with the design model of Flpp3-bound ASD1, with a Cα RMSD of 0.9 Å when aligned by Flpp3 residues and 0.4 Å when comparing the miniprotein alone (Figure 6A–B). The crystal structure confirms that ASD1 engages the α-site of Flpp3 through a network of hydrophobic and polar interactions, primarily mediated by the helices 2 and 3 of the miniprotein. The hydrophobic interactions at the core of the interface are mediated by ASD1 residues Leu25, Ala28, Leu33, Tyr44, Leu47, and Ala51, which pack against the complementary hydrophobic regions of the Flpp3 α-site. Additionally, several polar interactions contribute to the binding interface. ASD1 residues Leu25 and Tyr29 interact with Flpp3 residues Ser151 and Glu158, respectively (Figure 6C). Tyr44 and Asn39 of ASD1 are also optimally positioned to form hydrogen bonds with Gln88 on Flpp3 (Figure 6D). The binding interface is further stabilized by additional polar contacts involving ASD1 residues Arg52, Asp48, Tyr44, and Asn39, which interact with Flpp3 residues Arg149, Phe90, and Gln88, respectively (Figure 6E). Notably, the sidechain rotamers of interface residues in the X-ray crystal structure are nearly identical to those in the design model. Overall, the X-ray structure of ASD1-Flpp3 complex confirms the structure and binding mode of ASD1, highlighting the atomic accuracy provided by our computational design approach.

**Figure 6:**
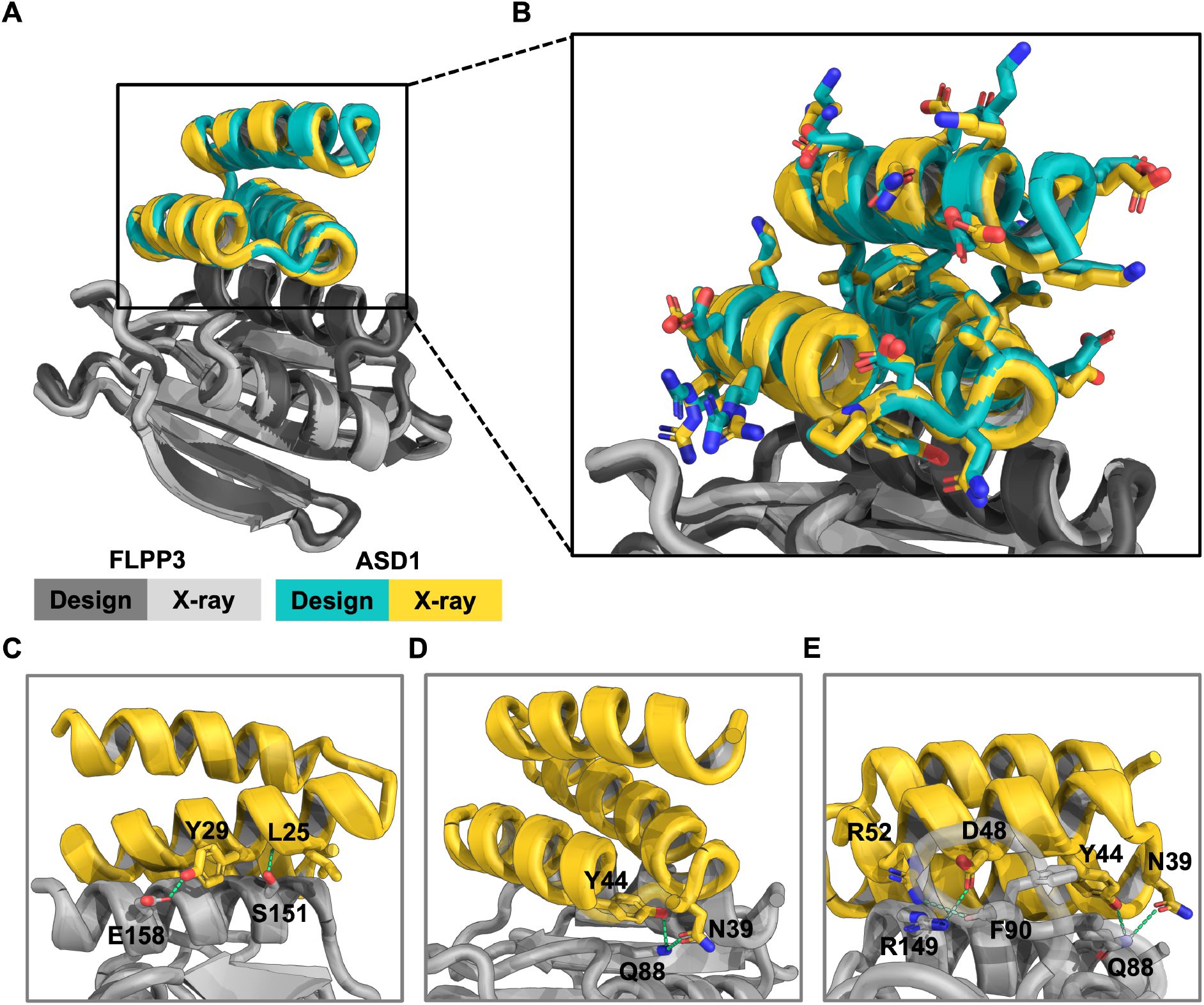
Crystal structure of the ASD1–Flpp3 complex confirms the designed binding mode. (A) Overall structure of the ASD1–Flpp3 complex. The Flpp3 crystal structure is shown in light gray, and ASD1 in yellow. The design models are overlaid in dark gray (Flpp3) and teal (ASD1). (B) Zoomed-in view of ASD1 design model and crystal structure with sidechains, showing close structural agreement. Cα RMSD for ASD1 is 0.9 Å when the structures are aligned by Flpp3 residues, and 0.4 Å when aligned by the binder only. (C–E) Key interactions between ASD1 and Flpp3, shown from different orientations of the interface. Key polar interactions between ASD1 and Flpp3 are shown as green dashed lines.

### Characterization of Flpp3 Surface Accessibility

To investigate the cellular location and orientation of Flpp3 in the outer membrane, we used flow cytometry to assess whether designed minibinders could label Flpp3 on the surface of intact *Francisella* cells. FITC-conjugated ASD1 minibinder was incubated with *Francisella novicida* at concentrations ranging from 0–10 μM. We observed no significant change in cell labeling between wild-type (WT) and Δ*flpp3* deletion strains at any of the tested concentrations (Figure S9B), suggesting that ASD1 does not bind Flpp3 on intact bacterial cells. To confirm if Flpp3 is expressed and potentially accessible on the surface of intact cells, we also generated a *F. novicida* strain expressing Flpp3 fused to a VSV-G tag. However, staining with anti-VSV-G antibody followed by a FITC-conjugated secondary antibody also showed no detectable staining compared to control strains (Figure S9A). Similar labeling studies with biotinylated β-site binder BSD1 and streptavidin-phycoerythrin (SAPE) at a range of concentrations (0 μM, 1 μM, 5 μM, and 10 μM) also showed no significant difference in fluorescence signal between the WT and knockout control strains (Figure S9C). Western blot analysis confirmed the expression of Flpp3 in the bacterial strains used for flow cytometry experiments (Figure S10), indicating that the lack of binding signal in intact cells was not due to expression issues. These results suggest that Flpp3 is either not accessible on the cell surface or is oriented in a manner that prevents minibinder access to either of the two selected sites. The complex lipopolysaccharides (LPS) barrier and outer capsule of Francisella may prevent access of medium-to-large molecular weight molecules, such as miniproteins and antibodies^7,23^. Alternatively, Flpp3 may be localized to the inner leaflet of the outer membrane or sequestered within oligomeric complexes, as has been proposed for other Francisella lipoproteins^24,25^, thereby rendering the selected binding sites inaccessible in intact cells.

## DISCUSSION

The natural and deliberate spread of *Francisella tularensis* remains a significant concern because of its exceptionally high infectivity and low doses required to cause a severe infection. In this work, we used a combination of physics-based and deep-learning computational methods to design high-affinity miniprotein binders targeting Flpp3, a key virulence factor of *F. tularensis*.

The computational approach described here overcomes several challenges in targeting Flpp3. Although the role of Flpp3 as a virulence factor is well established, structural and mechanistic understanding of Flpp3 and its interacting partners remains limited. The lack of Flpp3-ligand complex structures and well-defined binding pockets on Flpp3 surface make ligand discovery challenging. Despite this limited structural information, our computational methods enabled exploration of a vast conformational space to design and validate multiple miniprotein binders against two different sites of Flpp3. For site I, we identified four designed binders with nanomolar binding affinities, with the best binder displaying a K_D_ of 24 nM without any additional sequence optimization. While the initial binder for site II had a binding affinity of 81 nM, a round of site saturation mutagenesis identified multiple variants with binding affinities below 1 nM, including the best binder with a binding affinity of 580 pM. Circular dichroism spectroscopy confirmed that these binders fold into their designed all-helical topology. X-ray crystal structure of the best α-site binder, ASD1, in a complex with Flpp3 confirms that ASD1 engages Flpp3 through the designed interface, with close agreement between the experimental structure and the design model. The close agreement between the X-ray crystal structure and computational model of ASD1-Flpp3 complex (Cα RMSD: 0.9 Å) highlights the remarkable accuracy of our design methods and precision with which computationally designed miniproteins can be designed *de novo* to target difficult protein targets.

This work highlights the potential for computational approaches to custom design miniproteins with diverse shapes and sizes that bind bacterial virulence factors with very high affinity. The ability to computationally design protein binders with atomic level accuracy offers a powerful alternative to conventional library-based screening methods or antibodies, as it provides faster routes to affinity reagents with desired biophysical properties. The designed binders targeting different sites on Flpp3 also offer a means to precisely perturb its interactions with other proteins and explore the molecular mechanisms employed by *F. tularensis* during infection and immune evasion. While the flow cytometry experiments did not detect binder labeling on the surface of intact cells, these results suggest that Flpp3 may be positioned on the inner leaflet of the outer membrane or embedded within larger protein assemblies. In this context, the designed binders with their high specificity and affinity could serve as useful molecular tools to investigate Flpp3 localization, orientation, and structural environment in the native membrane using established tools like immunoprecipitation or fluorescence microscopy.

These miniproteins can further guide the development of small molecules and peptides with better access to the outer membrane proteins like Flpp3, penetration across the LPS and outer membrane, and superior drug-like properties. Future research efforts will focus on optimizing the selected binders, leveraging them to explore the precise mechanistic roles of Flpp3 in infection and immune evasion, and evaluating their therapeutic potential.

## METHODS

### *De novo* computational design

Miniprotein binders against Flpp3 were designed using a multi-step computational pipeline integrating physics-based docking and deep-learning-based sequence optimization. The soluble domain structure of Flpp3 (PDB ID: 6PNY) was used as the design target, and two distinct regions on it were selected to guide the binder design calculations: the α-site (α-helical, electropositive face) and the β-site (β-sheet, electronegative face). A hierarchical docking approach was applied to identify scaffold backbones with high binding potential. First, disembodied amino acid side chains were docked against the selected target regions using Rosetta RIFGen^13^ approach described previously. This was followed by global shape complementarity docking using PatchDock^17^, generating candidate scaffold placements. The docking results were refined using RIFDock^13^, which optimized rigid-body orientations and side-chain packing. A total of 500,000 docked conformations were generated for further sequence design. Sequence design was performed using two iterative rounds of ProteinMPNN^18^ and Rosetta FastRelax^26^ protocol for each miniprotein downselected from the Rosetta RIFDock steps. Designed sequences were filtered using Rosetta interface energy metrics (ddG < −40 kcal/mol for α-site, < −35 kcal/mol for β-site) and structural confidence scores from AlphaFold2 (pLDDT > 90 for α-site, > 80 for β-site; iPAE < 6.0)^15^. To remove aggregation-prone sequences, designs were further screened using spatial aggregation propensity (SAP) scores (<30 for α-site, <35 for β-site)^13^. The final selection yielded 15,000 α-site designs and 8,817 β-site designs, which were experimentally validated using yeast surface display and biochemical assays.

### Yeast surface display and fluorescence-activated cell sorting

We used yeast surface display (YSD) to screen and enrich miniprotein binders against Flpp3. Saccharomyces cerevisiae EBY100 cells were transformed with synthetic genes encoding the designed minibinders using a pETCON3 expression vector, enabling surface display via the Aga2p system^27^. Transformed yeast cells were cultured in C-Trp-Ura medium supplemented with 2% (w/v) glucose at 30°C, shaking at 220 rpm. For induction, cells were centrifuged, resuspended in SGCAA medium supplemented with 0.2% (w/v) glucose, and incubated at 30°C for 16–18 hours. The screening process consisted of four sequential fluorescence-activated cell sorting (FACS) rounds. The first round (called expression sorting) was performed without Flpp3 to isolate yeast cells successfully expressing minibinders on the surface. In the second round (initial avidity sorting), yeast cells were incubated with 1 μM Flpp3, anti-c-Myc fluorescein isothiocyanate (FITC) and streptavidin-phycoerythrin (SAPE) to enrich for binders. The third round, enrichment sorting, was conducted again at 1 μM Flpp3 to further increase the population of Flpp3-binding cells. In the final round (titration sorting), yeast cells were incubated with decreasing concentrations of Flpp3 (1,000 nM, 100 nM, 10 nM, 1 nM for the α-site library; 1,000 nM, 300 nM, 100 nM, 30 nM for the β-site library) to select for higher-affinity binders. FACS was performed using the Sony SH800 Cell Sorter, with gating thresholds established using a negative control population (yeast cells incubated without Flpp3) to define background fluorescence. Yeast populations were sorted based on this gated threshold, ensuring that only Flpp3-binding cells above background fluorescence were collected. Data were plotted and analyzed using FlowJo v10^28^. Sorted yeast populations were lysed using Zymolyase-Yeast Lytic Enzyme from the Zymoprep Yeast Plasmid Miniprep I kit, and enriched sequences were recovered for next-generation sequencing (NGS) to evaluate sequence enrichment across sorting rounds. Next-generation sequencing was performed following the protocol described by Cao et al. (2022)^13^.

### Flpp3 expression and purification

The gene encoding Flpp3, containing an N-terminal His6-Avi tag, was cloned into the pET-28a(+) expression vector and transformed into chemically competent E. coli BL21 (DE3) cells (NEB C2527I) following the recommended protocols. The plasmid contains a kanamycin resistance gene for selection. A single colony was inoculated into 3–4 mL of LB medium supplemented with kanamycin (50 µg/mL) and grown overnight at 37°C with shaking at 200 rpm. The overnight culture was then diluted into 1 L of 2XYT medium in 2 L baffled flasks and incubated at 37°C until reaching an OD600 of approximately 0.4. The incubation temperature was then reduced to 18°C, and after 30 min of acclimatization, protein expression was induced by adding isopropyl β-D-1-thiogalactopyranoside (IPTG) to a final concentration of 0.3 mM. The culture was incubated for an additional 16–20 hours at 18°C with continuous shaking. Cells were harvested by centrifugation at 9,000 × g for 20–30 min at 4°C, and the resulting cell pellet was either flash-frozen in liquid nitrogen for storage at −80°C or immediately processed for purification. For protein purification, the cell pellet was resuspended in 40 mL of lysis buffer per liter of culture (50 mM Tris-HCl, pH 7.5, 500 mM NaCl, 10% glycerol, 5 mM imidazole) supplemented with lysozyme (0.5 mg/mL), protease inhibitors (Pierce™ Protease Inhibitor, A32963), and Benzonase nuclease (Thermo Scientific, E1014) following manufacturer’s recommendations. The resuspended cells were lysed by sonication (30 s pulses at 50% amplitude, repeated for 4–5 cycles with 5–10 min cooling intervals on ice between rounds). The lysate was cleared by centrifugation at 17,000 × g for 30 min at 4°C, and the supernatant was transferred to a fresh tube. Purification was performed using immobilized metal affinity chromatography (IMAC) on a 1 mL HisTrap column (Cytiva) connected to an ÄKTA FPLC system. The column was equilibrated with Ni-Buffer A (50 mM Tris-HCl, pH 7.5, 500 mM NaCl, 10% glycerol, 5 mM imidazole), and the clarified lysate was loaded onto the column at a flow rate of 1 mL/min. The column was washed extensively with Ni-Buffer A to remove non-specifically bound proteins, and the target protein was eluted using an imidazole gradient (0–100% Ni-Buffer B containing 50 mM Tris-HCl, pH 7.5, 500 mM NaCl, 10% glycerol, and 500 mM imidazole). The eluted fractions were analyzed using SDS-PAGE, and those containing Flpp3 were pooled and concentrated using Amicon Ultra centrifugal filters (3 kDa MWCO) by centrifugation at 2,000–3,000 × g at 4°C. For further purification and buffer exchange, the protein was subjected to size exclusion chromatography (SEC) using a HiLoad 16/600 Superdex 200 column pre-equilibrated with sizing buffer (50 mM Tris-HCl, pH 7.5, 500 mM NaCl, 10% glycerol). The protein was loaded manually using a 1-2 mL sample loop and eluted at a flow rate of 0.5 mL/min. Fractions corresponding to Flpp3 were identified by A280 absorbance, pooled, and concentrated as described above. Protein concentration was determined using a NanoDrop spectrophotometer (Thermo Scientific) based on its predicted extinction coefficient. The purified protein was aliquoted, snap-frozen in liquid nitrogen, and stored at −80°C until further use.

### Miniprotein expression and purification

Minibinders were expressed in *E. coli* using a small-scale expression system in a 96-well deep-well plate format. Genes encoding the designed minibinders were synthesized by Integrated DNA Technologies (IDT) and cloned into the LM0627 (Addgene 191551) BVN2 expression vector with a C-terminal SNAC^29^ and 6×His tag for affinity purification. The plasmids were transformed into E. coli BL21(DE3) competent cells (New England Biolabs). Transformed cells were grown overnight in LB medium supplemented with kanamycin (50 µg/mL) at 37°C with shaking at 900 rpm in a deep-well plate incubator. The following day, overnight cultures were diluted 1:20 into fresh Terrific Broth (TB2) with autoinduction media supplemented with kanamycin (50 µg/mL) and 0.5% glycerol. Cultures were grown at 37°C with shaking for 18 h. Cells were harvested by centrifugation at 3,220 × g for 10 minutes at 4°C, and the resulting cell pellets were resuspended in 200 µL of lysis buffer per well. The lysis buffer consisted of BPER supplemented with 0.1 mg/ml lysozyme, 10 µg/ml DNase I and 1 mM PMSF. Lysis was carried out by incubation at room temperature with shaking for 30 minutes. The lysates were clarified by centrifugation at 4000 × g for 10 minutes at 4°C, and the supernatants were collected for purification. Minibinders were purified using Ni-NTA resin (Qiagen) in a high-throughput format. The resin was equilibrated with the wash buffer (50 mM Tris-HCl, 300 mM NaCl, 25 mM imidazole, pH 8.0) before being added to the clarified lysates. Binding was performed at room temperature for 1 hour, after which the resin was washed three times with wash buffer to remove unbound proteins. Minibinders were eluted in elution buffer (50 mM Tris-HCl, 300 mM NaCl, 500 mM imidazole, pH 8.0), and protein-containing fractions were collected. To further purify the minibinders, the eluates from Ni-NTA purification were subjected to size-exclusion chromatography (SEC) using a Superdex 75 Increase 10/300 GL column (Cytiva) equilibrated with SEC buffer (50 mM Tris-HCl, 150 mM NaCl, pH 8.0). Fractions corresponding to the expected molecular weight were collected and analyzed. Protein concentrations were determined using a NanoDrop spectrophotometer (Thermo Scientific) by measuring absorbance at 280 nm.

### Binding affinity determination using Biolayer Interferometry and Surface Plasmon Resonance

Binding affinities of the miniprotein hits identified from the yeast display screening were determined using biolayer interferometry using an Octet RED96 instrument (ForteBio). Streptavidin-coated biosensors (ForteBio) were first incubated with biotinylated Flpp3 at a concentration of 50–100 nM in BLI assay buffer (10 mM HEPES, 150 mM NaCl, 3 mM EDTA, 0.05% surfactant P20, and 1% BSA). Following target immobilization, biosensors were transferred to buffer-only wells to establish a baseline signal before being exposed to varying concentrations of minibinders to monitor the association phase. Next, the biosensors were returned to buffer-only wells for the dissociation phase to measure the off-rate (k_off_). The kinetic parameters, including association rate constant (k_on_), dissociation rate constant (k_off_), and equilibrium dissociation constant (K_D_), were determined using Octet Data Analysis software by fitting the data to a 1:1 binding model.

SPR experiments were conducted using a Cytiva Biacore 8K system with HBS-EP+ buffer (Cytiva) as the running buffer. Biotinylated Flpp3 was immobilized on a streptavidin-coated sensor chip using the Biotin Capture Kit (Cytiva). Prior to analyte injections, a capture test was performed to determine the optimal Flpp3 loading concentration, ensuring an appropriate surface density relative to the minibinders based on their molecular weight ratios. Binding interactions were assessed using single-cycle kinetics (SCK) at a flow rate of 30 µl/min. For the initial screening, minibinders were injected in a 4-point, 10-fold dilution series, with an association phase of 60 seconds followed by a 120-second dissociation phase. Based on the estimated K_D_ from the screening, a finer kinetic analysis was performed using a 4-point, 5-fold dilution series, extending the dissociation time to 500 seconds for improved resolution of slower off-rates. Sensorgrams were processed using Biacore Insight Evaluation Software, with double referencing applied. Binding kinetics were analyzed using a 1:1 binding kinetics fit model.

### Cell surface labeling

For bacterial flow cytometry, *F. novicida* U112 cells were cultured overnight in tryptic soy broth supplemented with 0.1% (w/v) cysteine (Research Products International C81020) (TSB-C) from a single colony at 37 °C with shaking. On the day of the experiment, bacteria were diluted 1:50 in TSB-C and subcultured to mid-log phase (OD600nm∼0.5). Bacteria were washed with sterile-filtered PBS with 1% bovine serum albumin (BSA; Research Products International A30075) and resuspended at approximately 1×10^7^ bacteria/mL. Minibinder was diluted in sterile PBS at various concentrations, and 25 µl of this solution was mixed with 25 µL diluted bacteria in a v-bottom plate. Cells were stained with minibinder for 1 hr at room temperature. For binding experiments using biotinylated minibinders, secondary staining of cells was performed with streptavidin-PE (Thermo Fisher 12-4317-87). Primary antibody staining of VSV-G tagged FLPP3 bacterial strains was performed with rabbit anti-VSV-G (Sigma Aldrich V4888), for 30 min at 4 °C. Cells were washed and stained with fluorophore-labeled secondary antibody (goat anti-rabbit-488; Invitrogen A-11008). Cells were washed and fixed in 4% paraformaldehyde (PFA). After washing, bacteria were resuspended in PBS and analyzed by FACS. Data were acquired on an LSR II (BD Biosciences).

### Bacterial strains and growth conditions

Bacterial strains used in this study include *F. tularensis* subspecies *novicida* U112 (*F. novicida*, gift from Colin Manoil, University of Washington, Seattle, WA), *Escherichia coli* strain DH5α (*E. coli* DH5α, Thermo Fisher Scientific), and *E. coli* BL21 (DE3). *F. novicida* strains were grown aerobically at 37 °C in tryptic soy broth or agar supplemented with 0.1% (w/v) cysteine (TSBC or TSAC). For selection, kanamycin was used at the following concentrations: 15 μg/mL (*F. novicida*) or 50 μg/mL (*E. coli*). *F. novicida* strains were stored in TSBC supplemented with 20% (v/v) glycerol at −80 °C. *E. coli* strains were stored in LB supplemented with 15% (v/v) glycerol at −80°C.

### Strain and plasmid construction

*F. novicida* Δ*flpp3* and *flpp3*–VSV-G strains were generated via allelic exchange as described previously^30–32^. Briefly, sequences containing 900 bp flanking the site of deletion or insertion were amplified by PCR and cloned into the BamHI and PstI sites of the vector pEX18-pheS-km using Gibson assembly^30^. Naturally competent *F. novicida* was prepared by back-diluting overnight cultures in TSBC, growing for 3 hrs at 37°C with shaking, harvesting by centrifugation, and resuspending in *Francisella* transformation buffer^31^. pEX18-pheS-km-based deletion or insertion plasmid was added to freshly prepared *F. novicida* competent cells. Bacterial suspensions were then incubated at 37 °C with shaking for 30 min, followed by addition of TSBC and an additional 3 hrs of incubation. Transformants were selected by plating on TSAC with kanamycin. The resulting merodiploids were grown overnight in non-selective TSBC, diluted into Chamberlain’s defined medium^33^ containing 0.1% p-chlorophenylalanine (w/v) and allowed to grow to stationary phase. Cultures were then streaked onto TSAC, colonies were patched onto TSAC with and without kanamycin, and kanamycin-sensitive colonies were screened for mutations by colony PCR. Primers used in plasmid and strain construction can be found in Supplementary Table 1.

### Flpp3–VSV-G expression analysis

To analyze the expression of Flpp3, *F. novicida* wild-type and *flpp3*–VSV-G strains were grown overnight in TSBC at 37 °C with shaking, back-diluted the next morning, grown to OD_600_∼0.9 and the equivalent of 1 ml culture at OD600=1 was collected for each strain by centrifugation. Cell pellets were resuspended in an equal volume of 1x Laemmli buffer^34^ and heated at 95 °C. Proteins in each sample were separated by SDS-PAGE and analyzed by western blotting as described previously^32^.

### Antibodies

Anti-VSV-G antibody produced in rabbit, Sigma V4888.

Anti-rabbit IgG (whole molecule)–peroxidase antibody produced in goat, Sigma A6154

### X-ray crystallography

All crystallization experiments were conducted using the sitting-drop vapor-diffusion method. Crystallization trials were set up in 200 nL drops using the 96-well plate format at 20°C. Crystallization plates were set up using a Mosquito LCP from SPT Labtech, then imaged using UVEX microscopes and UVEX PS-256 from JAN Scientific. Diffraction quality crystals formed in 0.1 M Bis-Tris Propane pH 6.5, 0.2 M Sodium iodide, 20% (w/v) PEG 3350 and 10% (v/v) Ethylene glycol for the Flpp3-ASD1 complex. Diffraction data were collected at the National Synchrotron Light Source II beamline FMX (17-ID-2). X-ray intensities and data reduction were evaluated and integrated using XDS^35^ and merged/scaled using Pointless/Aimless in the CCP4 program suite^36^. Structure determination and refinement starting phases were obtained by molecular replacement using Phaser^37^ using the designed model for the binder and 6PNY for the Flpp3 structure. Following molecular replacement, the models were improved using phenix.autobuild^38^; efforts were made to reduce model bias by setting rebuild-in-place to false, and using simulated annealing and prime-and-switch phasing. Structures were refined in Phenix^38^. Model building was performed using COOT^39^. The final model was evaluated using MolProbity^40^. Data collection and refinement statistics are recorded in Supplementary Table 2. Data deposition, atomic coordinates, and structure factors reported in this paper have been deposited in the Protein Data Bank (PDB), http://www.rcsb.org with accession code 9NLT.

### Mass spectrometry

To confirm the molecular mass of each miniprotein and expressed proteins, intact mass spectra were obtained via reverse-phase LC/MS on an Agilent G6230B TOF using an AdvanceBio RP-Desalting column.

### Plotting and Figures

Plots in this manuscript were generated using matplotlib^41^, seaborn^42^, FlowJo (v10), or GraphPad Prism (v10.4.1). Figures were created using PowerPoint v16.97.2 and BioRender. Protein structures were generated using PyMOL (version 2.5.2).

## Supporting information

Supplementary Information - Computational protocols and scripts

## ACKNOWLEDGEMENTS

We thank Lynda Stuart, Lance Stewart, Kandise VanWormer, Luki Goldschmidt, Kris Lindenauer, Sheharbano Jafry, Jonathan Palmer, and Katelyn Campbell for their helpful feedback, guidance, and support. We also thank the IPD Peptide, Crystallography, and Biologics, Vaccines, and Process Development (BioVaxPD) core labs, the UW Chemistry Mass Spectrometry Facility, and the Flow Cytometry Core at the UW Cell Analysis Facility Shared Resource Lab for providing instrumentation support and expertise. GG acknowledges support from the Fulbright Foreign Student Program during their PhD studies. This work was supported by funds from the DARPA Harnessing Enzymatic Activity for Lifesaving Remedies (HEALR) program HR001120S0052 contract HR0011–21–2–0012 (GB, DB, JDM, ML, LSK, JJW), start-up funds from the University of Washington’s Department of Medicinal Chemistry and Institute for Protein Design (GB), the Audacious Project (GB, AKB, AK), Marie Skłodowska–Curie Actions grant no. 101059124 (YFB), an NIH R01 grant no. R0AI160052 (AKB, AK). Crystallographic data were collected on NSLS II beamline 17-ID-2. The Center for Bio-Molecular Structure (CBMS) is primarily supported by the NIH-NIGMS through a Center Core P30 Grant (P30GM133893), and by the DOE Office of Biological and Environmental Research (KP1607011). NSLS-II is a U.S. DOE Office of Science User Facility operated under Contract No. DE-SC0012704.

## CONTRIBUTIONS

G.G.A., B.H., J.D.M., and G.B. conceived and conducted the study. G.G.A., G.B., L.S.K., Y.W., and A.K.B wrote the manuscript. G.G.A. and B.H. performed yeast surface display screenings. L.S.K. performed the bacterial cell binding flow cytometry assays. D.K.V., N.R., and I.G. carried out yeast library transformations. A.L. and S.R.G. expressed and purified Flpp3. V.A. and G.G.A expressed and purified the designed minibinders. A.L. and Y.F.B. prepared plasmids for Flpp3 expression. Y.W. designed and constructed the Δ*flpp3* and the *flpp3*–VSV-G strains, and performed the western blot experiment to confirm Flpp3 expression in the relevant strains. S.R. contributed to initial SPR assay development and performed preliminary experiments. G.G.A. conducted subsequent SPR experiments and data analysis. X.L. verified minibinder integrity via mass spectrometry. A.K.B. and A.K. determined the X-ray crystal structures of the designed minibinders bound to Flpp3. D.B., J.D.M., J.J.W., and G.B. supervised the project. All authors contributed to data interpretation and manuscript revision. G.G.A. and B.H. agree to be considered co-first authors and that their names may be reordered for personal or professional purposes.

## CODE AVAILABILITY

Computational protocols and scripts used for this work are available as Supplementary Files.

