## Supplementary Information - Figures and tables for "De novo design of high-affinity miniprotein binders targeting *Francisella tularensis* virulence factor"

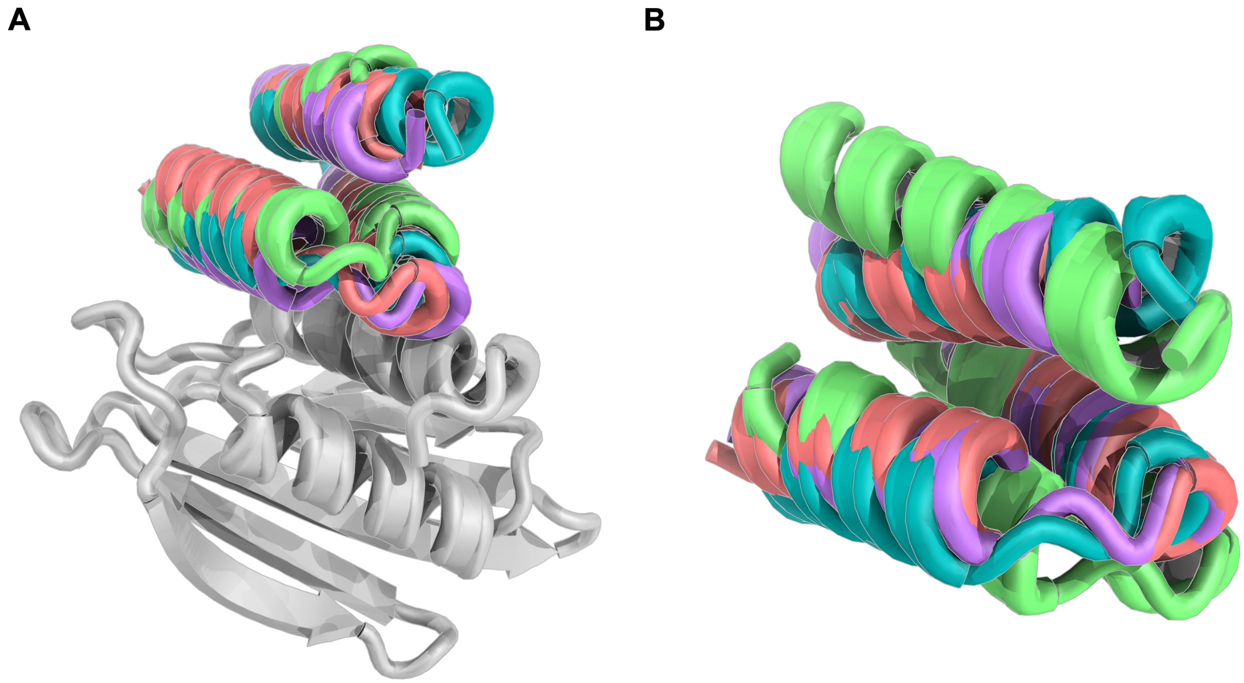


**Figure S1: Structural comparison of computationally designed α-site binders (ASD1–4).** (A) Design models of ASD1 (teal), ASD2 (salmon), ASD3 (green), and ASD4 (purple) are shown superimposed in their binding poses on Flpp3 (gray). All four binders adopt a similar three-helix fold and target the same region of Flpp3, with ASD1 and ASD4 displaying nearly identical backbone conformations and interface contacts. ASD2 engages the same epitope but with a modest shift in helix orientation, while ASD3 adopts a more divergent fold and binding geometry. (B) Structural alignment of ASD1–4 based on Cα RMSD highlights differences in backbone geometry independent of Flpp3 binding context. Alignments were performed in PyMOL using backbone (Cα) atoms only, without reference to the target.

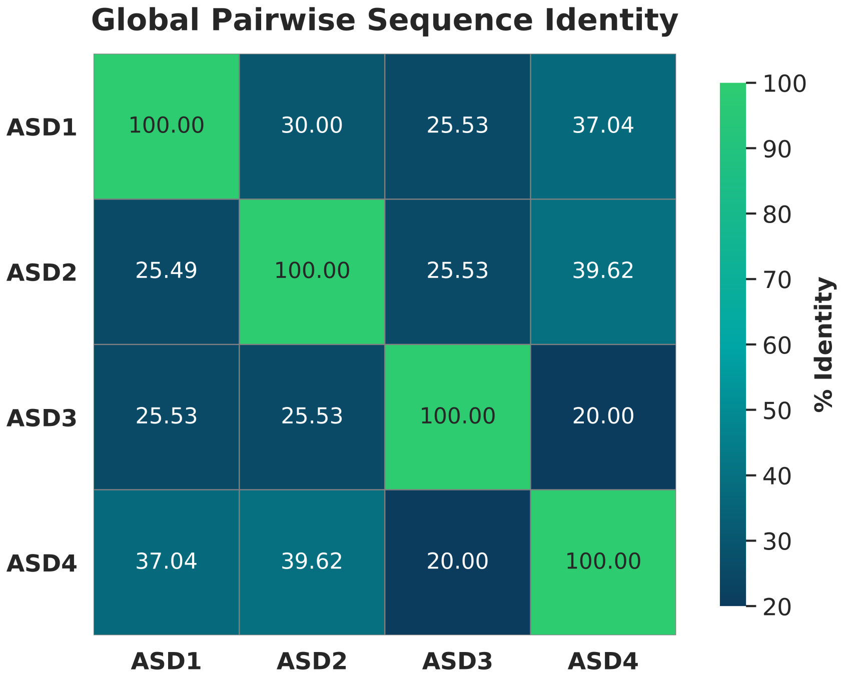


**Figure S2: Global pairwise sequence identity among α-site binders.**
Pairwise sequence identity heatmap for ASD1-ASD4 minibinders, calculated using global sequence alignment with the BLOSUM62 substitution matrix. Percent identities were computed based on aligned residues using the Needleman-Wunsch algorithm[44], with a gap opening penalty of –10 and a gap extension penalty of –1. Green indicates higher sequence identity, while darker blue corresponds to lower identity.

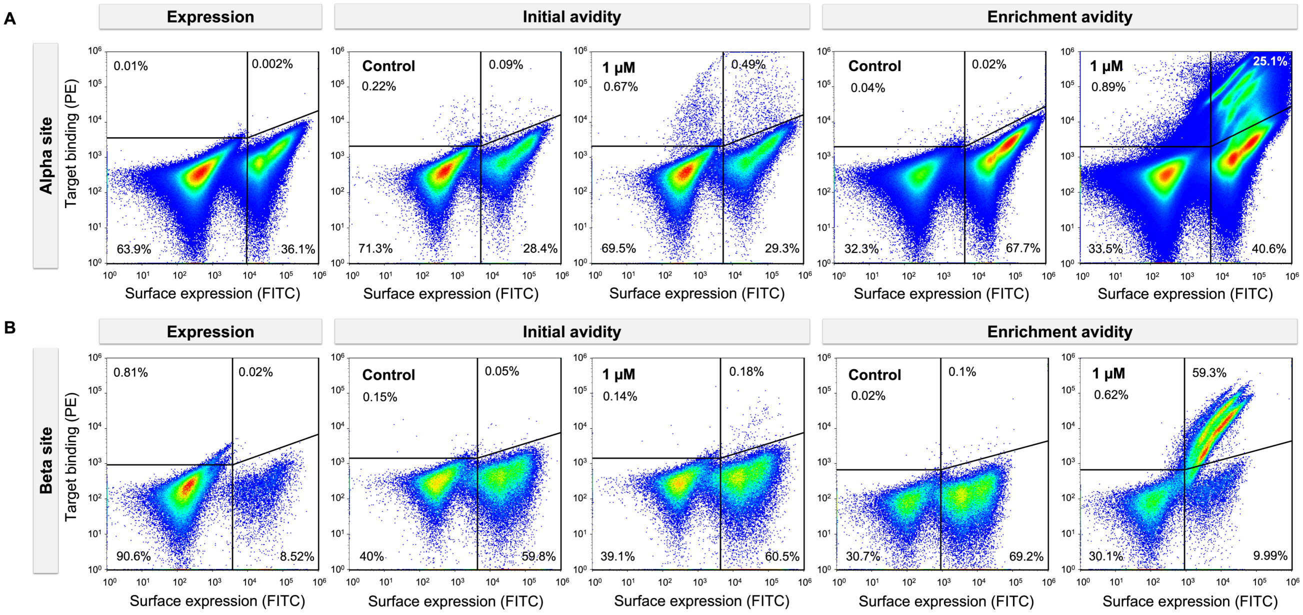


**Figure S3: Yeast surface display screening of α-site and β-site libraries against Flpp3 through expression and avidity-based enrichment.**
(A) Binding profiles of the α-site library during expression, initial avidity, and enrichment avidity sorts. A distinct population of Flpp3-binding cells is observed in the enrichment avidity sort at 1 μM. (B) Binding profiles of the β-site library under the same conditions. A clear binding population is also observed in the Enrichment avidity sort at 1 μM.

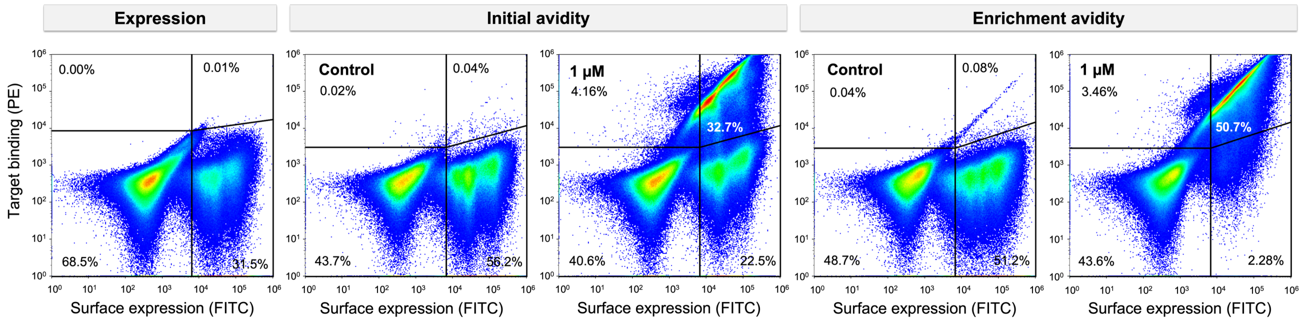


**Figure S4: Yeast surface display screening of BSD1 variants against Flpp3 through expression and avidity-based enrichment.**
Binding profiles of the BSD1 library during expression, initial avidity, and enrichment avidity sorts. A distinct population of Flpp3-binding cells is observed starting from the initial avidity sort at 1 μM, with further enrichment seen in the enrichment avidity sort.

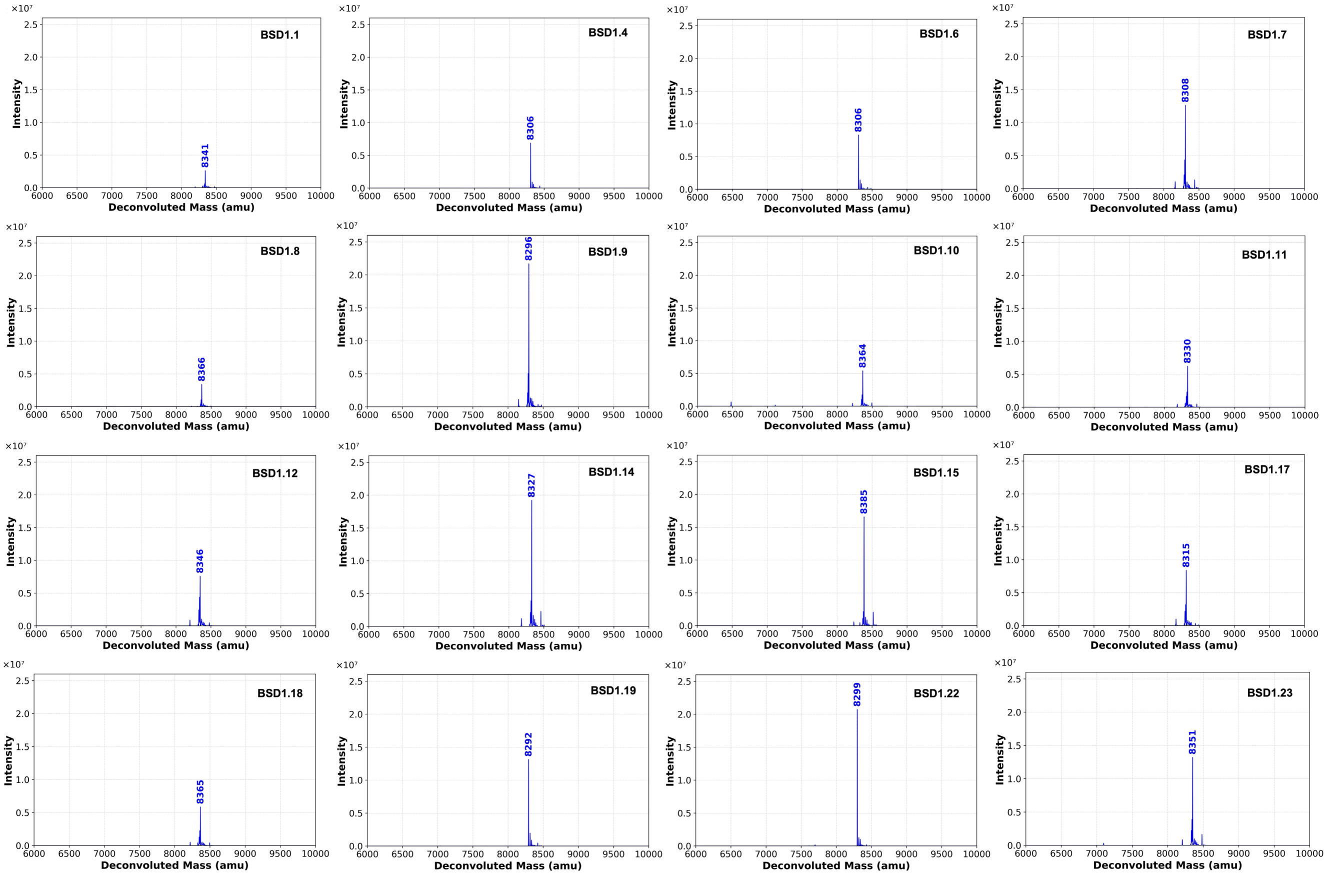


**Figure S5: Mass spectrometry analysis of expressed BSD1 variants.**

Deconvoluted electrospray ionization mass spectrometry (ESI-MS) spectra of selected BSD1 variants show single peaks corresponding to the expected molecular weights. All observed masses match the theoretical values, confirming successful expression and correct mass of the purified proteins.

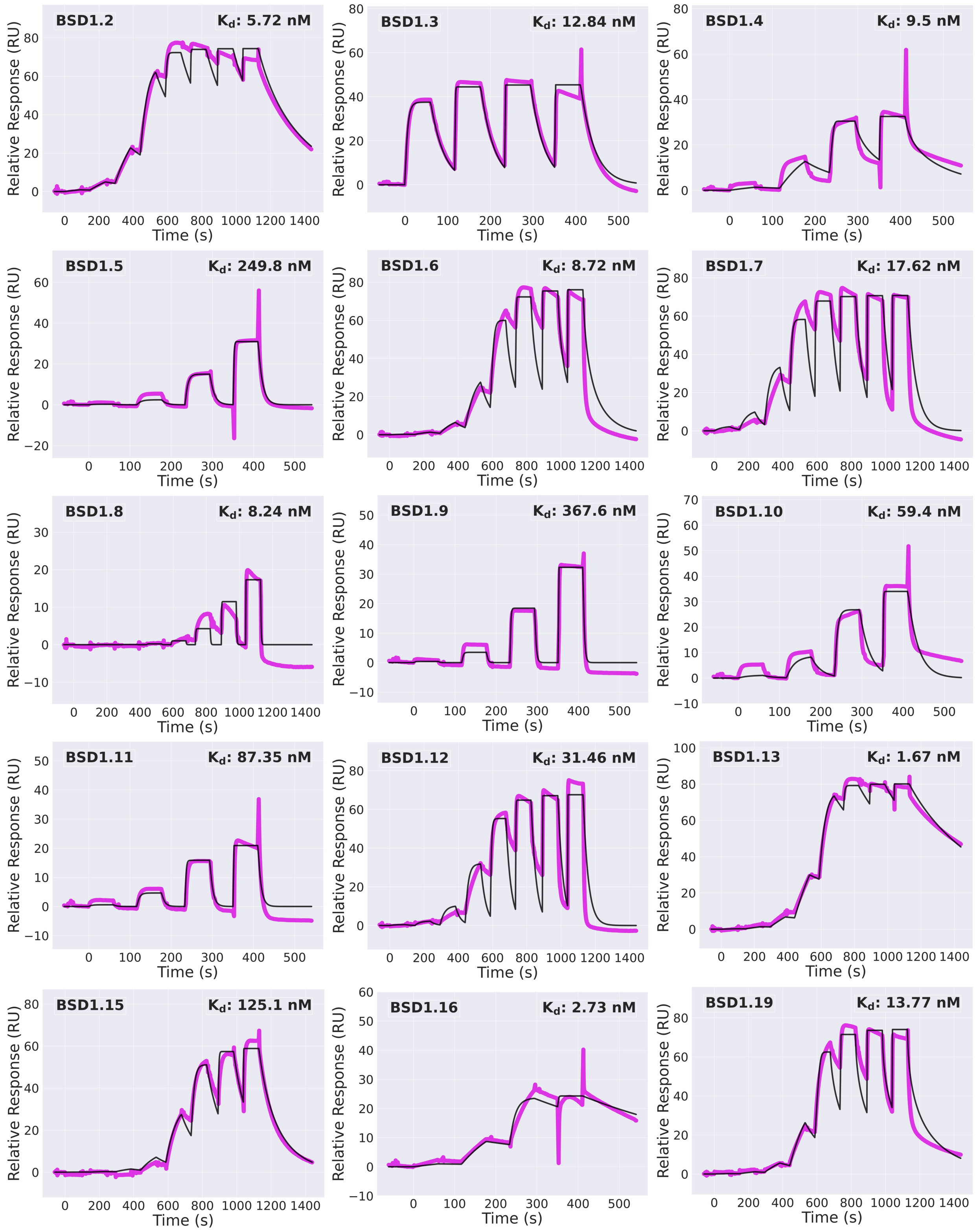


**Figure S6: SPR-based binding screen of additional BSD1 variants against Flpp3.**
SPR sensorgrams from a 4-point single cycle kinetics experiment (5-fold dilution, with highest concentrations ranging from 0.5 µM to 50 µM depending on the minibinder). Experimental data are shown in magenta, and global fits are shown with black lines.

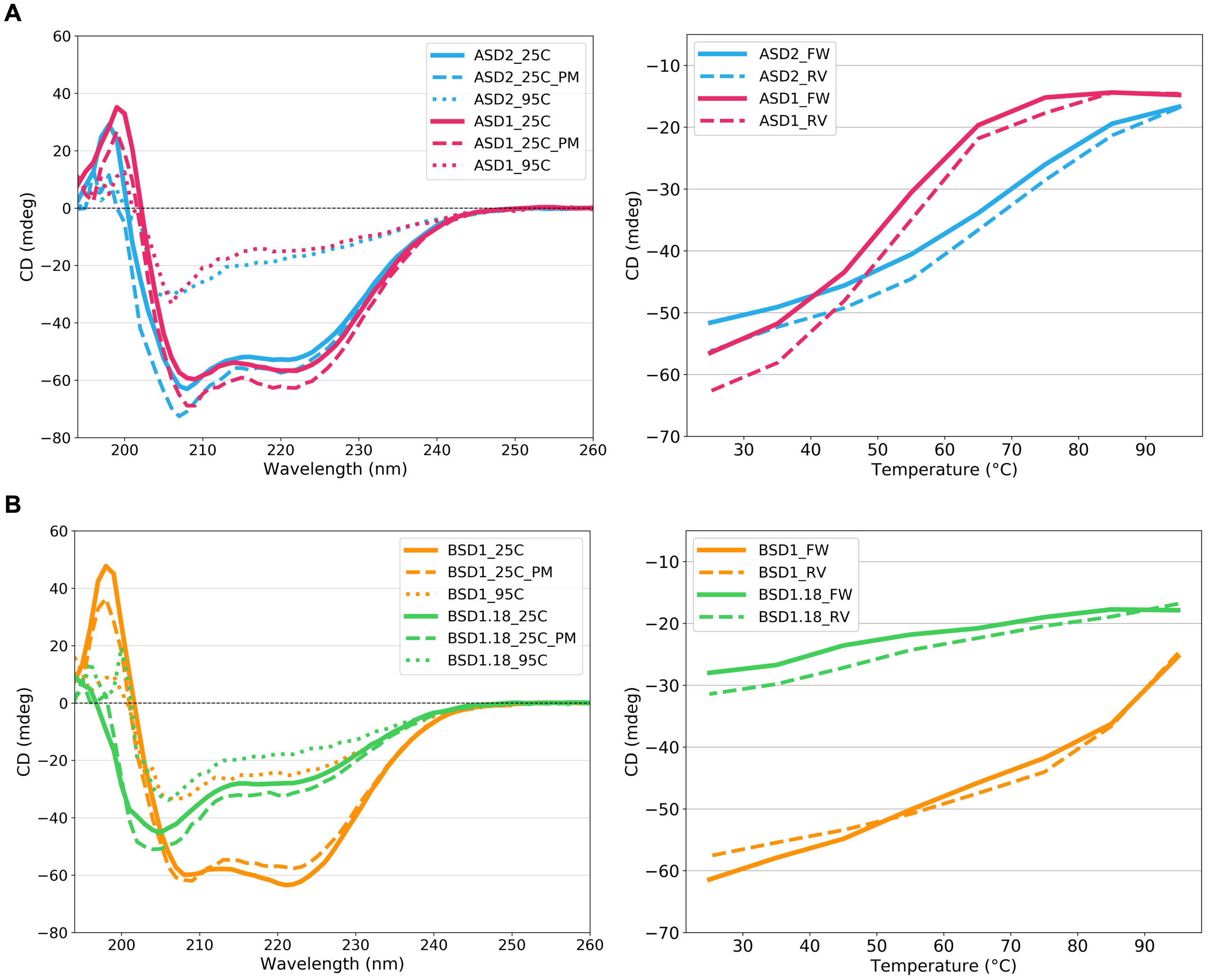


**Figure S7: Circular dichroism (CD) analysis of designed Flpp3 minibinders.**
CD spectra between 195–260 nm for the top α-site binders (panel A) and β-site binders (panel B). (Left) Spectra recorded at 25°C, 95°C, and after cooling back to 25°C (labeled as 25C_PM). (Right) Thermal melt curves showing changes in the CD signal at 222 nm as the temperature was increased to 95°C (solid lines, labeled _FW) and then cooled back to 25°C (dashed lines, labeled _RV).

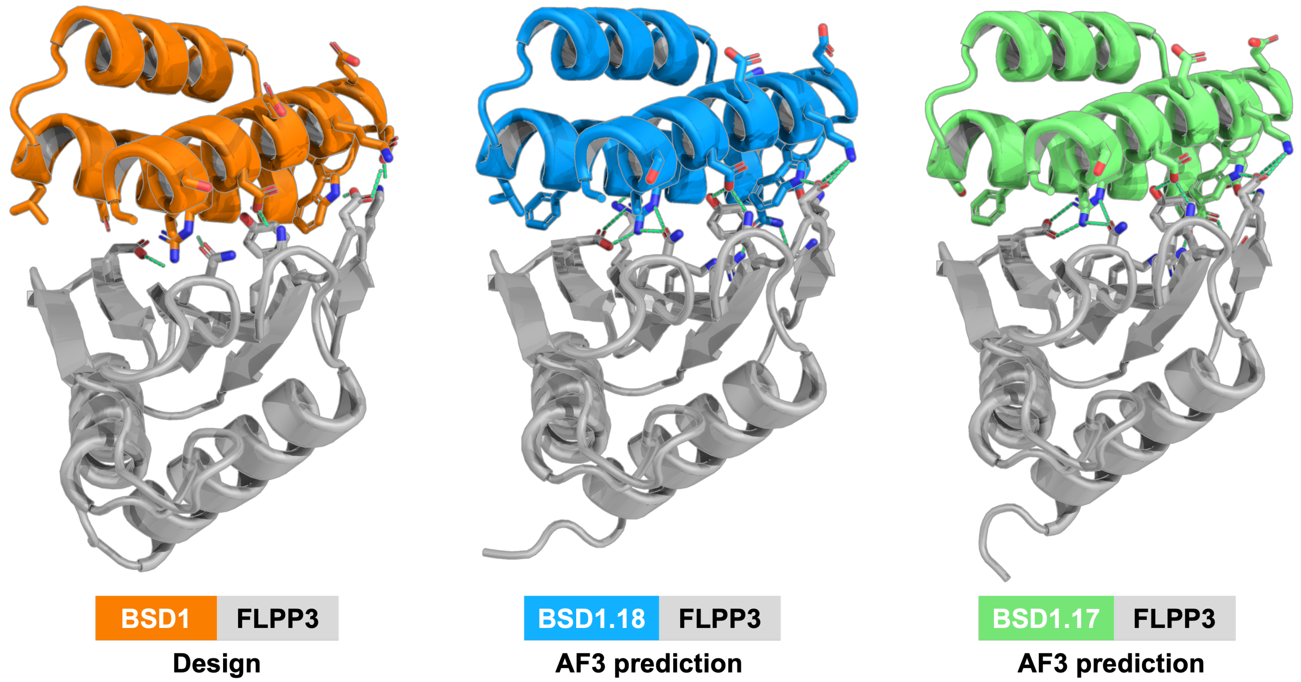


**Figure S8: Structural comparison of the original BSD1 design and AlphaFold3-predicted models of top binding variants.**

The de novo designed BSD1 miniprotein (left, orange) is shown in complex with Flpp3 (gray), alongside AlphaFold3-predicted structures of the two highest-affinity variants, BSD1.18 (middle, blue) and BSD1.17 (right, green). Both variants closely match the overall fold and binding mode of the original design, suggesting that the interface remains largely unchanged.

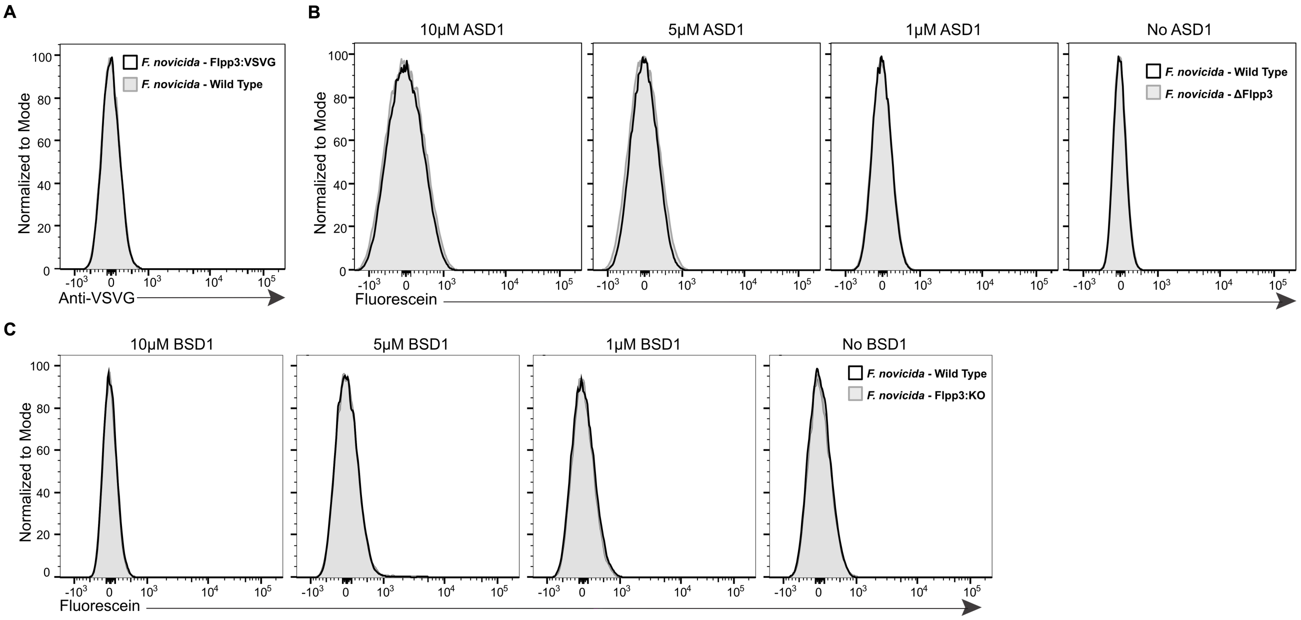


**Figure S9: Flow cytometry analysis of surface expression and minibinder binding in *F. novicida*.**

(A) *F. novicida* strains expressing Flpp3-VSV-G were stained with an anti-VSV-G antibody followed by a FITC-labeled secondary antibody to assess surface expression. Wild-type (WT) bacteria were included as a control. (B) WT and Δ*flpp3 F. novicida* strains were stained with FITC-labeled ASD1 minibinders at concentrations of 10 μM, 5 μM, 1 μM, and 0 μM. (C) WT and Flpp3 knockout (KO) strains were incubated with a biotinylated BSD1 minibinder and detected with streptavidin-phycoerythrin (SAPE). Minibinder concentrations were 10 μM, 5 μM, 1 μM, and 0 μM. Histograms represent fluorescence intensity normalized to mode.

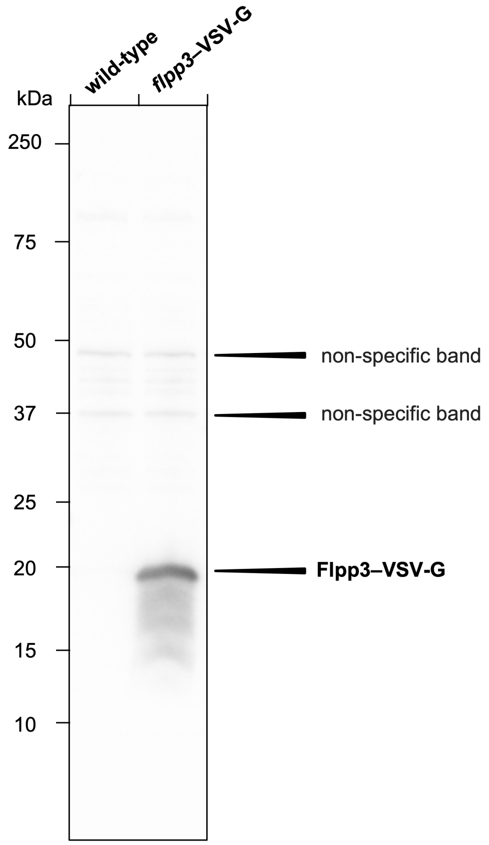


**Figure S10: Western blot analysis of *flpp3*–VSV-G expression in *F. novicida*.**
Whole-cell lysates from *F. novicida* wild-type and *flpp3*–VSV-G strains were probed with an anti–VSV-G antibody to detect expression of the Flpp3–VSV-G fusion protein. A strong band is observed in the *flpp3*–VSV-G lane, indicating successful expression. Molecular weights were estimated based on the Precision Plus Protein Standards Dual Color ladder (Bio-Rad Cat#1610374). Image shown is the chemiluminescent signal acquired using the iBright™ imaging system, with no contrast adjustments applied. Lane identities are labeled above the blot.

**Supplementary Table 1: Oligonucleotides used in this study.**

| Oligonucleotides or linear DNA fragments | Source |
|---|---|
| F1_ΔFTN_1382_BamHI (5' – 3') ATTCGAGCTCGGTACCCGGGGATCCTTGCTACGATGTTGTAATTGTTTATC | Integrated DNA Technologies |
| R1_ΔFTN_1382 (5' – 3') GAATATTTCCTTTCATATTATTTATTGATATACATAGTTCAG | Integrated DNA Technologies |
| F2_ΔFTN_1382 (5' – 3') TAATATGAAAGGAAATATTCGCTACGCTAATAC | Integrated DNA Technologies |
| R2_ΔFTN_1382_PstI (5' – 3') GCCAAGCTTGCATGCCTGCAGTGTTTCTGGTACTAGCAATTTTG | Integrated DNA Technologies |
| Check1_ΔFTN_1382 (5' – 3') CTGATAGAGTGCCTTGAGAATCC | Integrated DNA Technologies |
| Check2_ΔFTN_1382 (5' – 3') ATGTCAGATCTGCATATACTGTAGC | Integrated DNA Technologies |
| F1_FTN_1382_VSVG_BamHI (5' – 3') ATTCGAGCTCGGTACCCGGGGATCCTCGTATGTATAAACCTGTTTCTC | Integrated DNA Technologies |
| R1_FTN_1382_VSVG (5' – 3') TTTTCCTAATCTATTCATTTCAATATCTGTATATGTATTAGCGTAGCGAATATTTC | Integrated DNA Technologies |
| F2_FTN_1382_VSVG (5' – 3') TATTGAAATGAATAGATTAGGAAAATAATATGTAAAACTTAAAACTTTTATATATGATTG | Integrated DNA Technologies |
| R2_FTN_1382_VSVG_PstI (5' – 3') GCCAAGCTTGCATGCCTGCAGAACAATCTTCGAAGCAAAATATG | Integrated DNA Technologies |
| Check1_FTN_1382_VSVG (5' – 3') GCCAGCATGCGACATTGATAAG | Integrated DNA Technologies |
| Check2_FTN_1382_VSVG (5' – 3') TTTAAGTTTTACATATTATTTTCCTAATCTATTCATTTC | Integrated DNA Technologies |

*FTN_1382 corresponds to *flpp3*.

**Supplementary Table 2: Data collection and refinement statistics**

| | ***ASD1 - Flpp3 complex (PDB ID: 9NLT)*** |
|---|---|
| **Data collection** | |
| Space group | $P\ 2_1$ |
| Cell dimensions | |
| *a*, *b*, *c* (Å) | 36.17, 42.89, 54.05 |
| α, β, γ (°) | 90, 96.28, 90 |
| Resolution (Å) | 35.95 - 2.37 (2.50 - 2.37) |
| $R_{merge}$ | 0.327 (1.475) |
| $I / \sigma I$ | 4.3 (1.9) |
| Completeness (%) | 95.0 (99.9) |
| Redundancy | 6.3 (6.5) |
| **Refinement** | |
| Resolution (Å) | 35.95 - 2.37 (2.56 - 2.37) |
| No. reflections | 6458 (1233) |
| $R_{work}$ / $R_{free}$ | 0.2218 (0.3330) /0.2688 (0.3825) |
| No. atoms | |
| Protein | 1266 |
| Ligand/ion | 2 |
| Water | 11 |
| *B*-factors | |

| | |
|---|---|
| Protein | 51 |
| Ligand/ion | 29 |
| Water | 38 |
| R.m.s. deviations | |
| Bond lengths (Å) | 0.002 |
| Bond angles (°) | 0.460 |

*Single Crystal used for each data collection.
*Values in parentheses are for the highest-resolution shell.

**Supplementary Table 3: Amino acid sequences of the FIpp3 minibinders**

| ***Name*** | ***Sequence*** |
|---|---|
| ASD1 | MEEKEKEFNEKLEELKKAKTEEEKLELAYECGLLAGEINDPKYYRALDEVARAK |
| ASD2 | GREEVEKLCEEAVKEKDEKKREELIRRAAEIAAGYNDQESLKLVWDAIEKIES |
| ASD3 | SKEEEDVELVLKEIDKLVADGQPEIAKIVAEKVVEHLEELGNPDLAKRVRDKLEEI |
| ASD4 | EVMEKVKKLCEEAEEAKKAGNWEKVEELMRKAGLVAGEAGDLEACQLVDKKAKELEE |
| BSD1 | LISSVLQDRLETAKKWAEEGDKENAKFLLESAKQLAELVGDEETVKECEELLKKI |
| BSD1.1 | LISSVLQDRLETAKKWAEEGDKENAKFLLESAK**F**LAELVGDEETVKECEELLKKI |
| BSD1.2 | LISSVLQDRLETAKKWAEEGDKENAKFLLESAK**Y**LAELVGDEETVKECEELLKKI |
| BSD1.3 | LISSVLQDRLETAKKWAEEGDKENAKFLLESAK**V**LAELVGDEETVKECEELLKKI |
| BSD1.4 | LISSVLQDRL**L**TAKKWAEEGDKENAKFLLESAKQLAELVGDEETVKECEELLKKI |
| BSD1.5 | LISSVLQDRLETAKKWA**N**EGDKENAKFLLESAKQLAELVGDEETVKECEELLKKI |
| BSD1.6 | LISSVLQDRLETAKKWA**L**EGDKENAKFLLESAKQLAELVGDEETVKECEELLKKI |
| BSD1.7 | LISSVLQDRLETAKKWAEEGDKENAKFLL**D**SAKQLAELVGDEETVKECEELLKKI |
| BSD1.8 | LISSVLQDRLETAKKWAEEGDKENAKFLLES**D**KQLAELVGDEETVKECEELLKKI |
| BSD1.9 | LISSVLQDRLETAKKWAEEGDKENAKFLLESAKQLAE**S**VGDEETVKECEELLKKI |
| BSD1.10 | LISSVLQDRLETAKKWAEEGDKENAKFLLESAKQLAELV**V**DEETVKECEELLKKI |
| BSD1.11 | LISSVLQDRLETAKKWAEEGDKENAKFLLESAKQLAELVGD**H**ETVKECEELLKKI |
| BSD1.12 | LISSVLQDRLETAKKWAEEGDKENAKFLLESAKQLAELVGDEETVKECEELLKK**H** |
| BSD1.13 | LISSVLQDRL**L**TAKKWAEEGDKENAKFLLESAK**F**LAELVGDEETVKECEELLKKI |
| BSD1.14 | LISSVLQDRLETAKKWAEEGDKENAKFLL**D**SAK**F**LAELVGDEETVKECEELLKKI |
| BSD1.15 | LISSVLQDRLETAKKWAEEGDKENAKFLLES**D**K**F**LAELVGDEETVKECEELLKKI |
| BSD1.16 | LISSVLQDRLETAKKWAEEGDKENAKFLLESAK**F**LAELVGD**H**ETVKECEELLKKI |
| BSD1.17 | LISSVLQDRLETAKKWAEEGDKENAKFLLESAK**F**LAE**S**VGDEETVKECEELLKKI |
| BSD1.18 | LISSVLQDRLETAKKWAEEGDKENAKFLLESAK**F**LAELVGDEETVKECEELLKK**H** |
| BSD1.19 | LISSVLQDRL**L**TAKKWAEEGDKENAKFLL**D**SAKQLAELVGDEETVKECEELLKKI |

| | |
|---|---|
| BSD1.20 | LISSVLQDRL**L**TAKKWAEEGDKENAKFLL**D**SAK**F**LAELVGDEETVKECEELLKKI |
| BSD1.21 | LISSVLQDRL**L**TAKKWAEEGDKENAKFLLESAK**F**LAELVGD**H**ETVKECEELLKKI |
| BSD1.22 | LISSVLQDRL**L**TAKKWAEEGDKENAKFLLESAK**F**LAE**S**VGDEETVKECEELLKKI |
| BSD1.23 | LISSVLQDRLETAKKWAEEGDKENAKFLL**D**SAK**F**LAELVGDEETVKECEELLKK**H** |
| BSD1.24 | LISSVLQDRL**L**TAKKWAEEGDKENAKFLL**D**SAKQLAELVGD**H**ETVKECEELLKKI |
