## Supplementary Information - Computational protocols and scripts for "De novo design of high-affinity miniprotein binders targeting *Francisella tularensis* virulence factor": auto_relax_beta_v3.1.sh

```
#!/bin/bash
#SBATCH --mem=16g
#SBATCH -o auto_relax_beta_v3.1.log 
#SBATCH -e auto_relax_beta_v3.1.err 

set -x

###########################################---HOW TO RUN THIS SCRIPT---##################################################
#Make sure you check out an interactive node with enough memory. Example: qlogin -c 1 --mem=100gb
#Next type the following into your directory containing this script: chmod 755 auto_relax.sh
#Next, type: ./auto_relax.sh
#You will get a prompt asking you for PDB IDs. Just type the 4-letter/number and hit enter. Do NOT include .pdb 
#GOOD EXAMPLE INPUT: 1AB1 1AB2 1AB3
#BAD EXAMPLE INPUT: 1AB1.pdb 1AB2.pdb 1AB3.pdb
#########################################################################################################################


####################################---WHEN THIS SCRIPT WILL NOT WORK---#################################################
#If the .PDB file cannot be downloaded (too large). You will need to manually convert the PDBx/mmCIF or PDBML/XML to the standard PDB file format first.
#If the authors of the PDB entry did not desposit their structure factor information. Not sure what to do about this.
#########################################################################################################################

$SLURM_SUBMIT_DIR

ln -s /home/longxing/Rifdock/rosetta_master/source/src/apps/public/relax_w_allatom_cst/clean_pdb_keep_ligand.py ./step1_cleanPDB.py
ln -s /home/longxing/Rifdock/rosetta_master/source/src/apps/public/symmetry/make_symmdef_file.pl ./step2_make_xtalsym.pl 
source /software/phenix/setup.sh

arr=("$@")
if test -z "$arr"; then
	read -r -p "Enter PDB ID numbers you want to download and relax, separated by a single space: " -a arr
fi

################---This downloads your PDBs and formats your structure factor information using phenix---##############

for target in "${arr[@]}"; do
	echo "Downloading ${target}.pdb..."
	phenix.fetch_pdb ${target}
	echo "Downloading structure factor information for ${target}.pdb..."
	phenix.fetch_pdb -x ${target}
	echo "Converting reflection data for ${target}.pdb..."
	phenix.cif_as_mtz ${target}-sf.cif --merge --symmetry=${target}.pdb

	echo "Checking for intensity data to convert to amplitude data and generating a new set of R-free-flags for ${target}.pdb..."
	phenix.reflection_file_converter ${target}-sf.mtz --label="IOBS,SIGIOBS" --write_mtz_amplitudes --mtz_root_label=FOBS --generate_r_free_flags --mtz=${target}-sf_convert_rfree.mtz
	echo "Checking for amplitude data and generating a new set of R-free-flags for ${target}.pdb..."
	phenix.reflection_file_converter ${target}-sf.mtz --label="FOBS,SIGFOBS" --write_mtz_amplitudes --mtz_root_label=FOBS --generate_r_free_flags --mtz=${target}-sf_rfree.mtz

	if test -f "${target}-sf_rfree.mtz"; then
		echo "Amplitude data was found for ${target}.pdb."
		mv ${target}-sf_rfree.mtz ${target}-sf.mtz
		rm ${target}-sf_convert_rfree.mtz
	elif test -f "${target}-sf_convert_rfree.mtz"; then
		echo "Intensity data was converted to amplitude data for ${target}.pdb."
		mv ${target}-sf_convert_rfree.mtz ${target}-sf.mtz
		rm ${target}-sf_rfree.mtz
	fi 
done

#########################################################################################################################


######################---This section formats/cleans from your pdb file and creates symmetry info---#####################

for target in "${arr[@]}"; do

	echo "Removing non-protein atoms..."
	egrep -v "CONECT|HET" ${target}.pdb > tmp.pdb

	echo "Cleaning ${target}.pdb..."
	python2 step1_cleanPDB.py tmp.pdb -ignorechain
	mv tmp.pdb_00.pdb ${target}_clean.pdb
	rm tmp.pdb

	echo "Grabbing symmetry data and applying it to cleaned ${target}.pdb..."
	grep 'CRYST1' ${target}.pdb >> ${target}_clean.pdb

	echo "Creating symmetry file for ${target}.pdb"
	perl step2_make_xtalsym.pl -m cryst -p ${target}_clean.pdb > ${target}_cryst.symm

	source /home/longxing/phenix-1.11/phenix-1.11.1-2575_frank/phenix_env.sh

done

########################################################################################################################


##################################---This section runs the relax_into_density script---###################################

for target in "${arr[@]}"; do

	phenix.rosetta.run_phenix_interface /home/longxing/Rifdock/rosetta_rifdock/source/bin/rosetta_scripts.python.linuxgccrelease -database /home/longxing/Rifdock/rosetta_rifdock/database/ -parser:protocol /home/longxing/Rifdock/rosetta_rifdock/source/src/apps/public/crystal_refinement/low_resolution_refine.xml -parser::script_vars input.xray_data.r_free_flags.generate=True symmdef=${target}_cryst.symm bfactstrat=individual map_type=Auto -s ${target}_clean.pdb -mtzfile ${target}-sf.mtz -crystal_refine -set_weights cart_bonded 0.5 pro_close 0 -ignore_unrecognized_res

mv ${target}_clean_0001.pdb ${target}_ready.pdb

done

##########################################################################################################################
```
