## Supplementary Information - Computational protocols and scripts for "De novo design of high-affinity miniprotein binders targeting *Francisella tularensis* virulence factor": dl_interface_design_R2.py

```
#!/usr/bin/env python

import os, sys
import math

os.environ['XLA_PYTHON_CLIENT_PREALLOCATE']='false'

sys.path.insert( 0, '/net/software/pyrosetta3.10/custom/dev_nrbennet_py39_build1/build1' )

from pyrosetta import *
from pyrosetta.rosetta import *

import numpy as np
from collections import defaultdict
from collections import OrderedDict
import time
import argparse
import itertools
import subprocess
import time
#import pandas as pd
import glob
from decimal import Decimal

import torch

sys.path.append( '/home/nrbennet/software/dl/dl_design/mpnn/noise_mpnn/nate_scripts' )
import util_protein_mpnn as mpnn_util

sys.path.append( '/home/nrbennet/software/silent_tools' )
import silent_tools

init( "-beta_nov16 -in:file:silent_struct_type binary" +
    " -holes:dalphaball /work/tlinsky/Rosetta/main/source/external/DAlpahBall/DAlphaBall.macgcc" +
    " -use_terminal_residues true -mute basic.io.database core.scoring" )

def cmd(command, wait=True):
    the_command = subprocess.Popen(command, shell=True, stdout=subprocess.PIPE, stderr=subprocess.PIPE, universal_newlines=True)
    if (not wait):
        return
    the_stuff = the_command.communicate()
    return str( the_stuff[0]) + str(the_stuff[1] )

def range1( iterable ): return range( 1, iterable + 1 )

#################################
# Argument Validation Functions
#################################

def design_xml_check( input_val ):
    if not os.path.exists( input_val ):
        raise argparse.ArgumentTypeError( 'The argument -design_xml_check must be a path to an xml file, you used: %s'%input_val )
    return input_val

def feature_check( input_val ):
    if input_val not in [ 'full', 'coarse' ]:
        raise argparse.ArgumentTypeError( 'The argument -protein_features must be either full or coarse, you used: %s'%input_val )
    return input_val


#################################
# Parse Arguments
#################################

parser = argparse.ArgumentParser()
parser.add_argument( "-silent", type=str, default="", help='The name of a silent file to run this metric on. pdbs are not accepted at this point in time' )
parser.add_argument( "-checkpoint_path", type=str, default='/home/justas/projects/lab_github/proteinmpnn/model_weigths/pb58_020/epoch178_step265652.pt' )
parser.add_argument( "-relax_cycles", type=int, default="0", help="The number of MPNN->FastRelax cycles to perform (default 2)" )
parser.add_argument( "-temperature", type=float, default=0.1, help='An a3m file containing the MSA of your target' )
parser.add_argument( "-augment_eps", type=float, default=0.05, help='The variance of random noise to add to the atomic coordinates (default 0.05)' )
parser.add_argument( "-output_intermediates", action="store_true", help='Whether to write all intermediate sequences from the relax cycles to disc (defaut False)' )
parser.add_argument( "-do_predictor", action="store_true", help='Whether or not to run the predictor step and generate an estimated ddg for each MPNN sequence (default False)' )
parser.add_argument( "-protein_features", type=feature_check, default='full', help='An a3m file containing the MSA of your target' )
parser.add_argument( "-omit_AAs", type=str, default='X', help='A string off all residue types (one letter case-insensitive) that you would not like to use for design. Letters not corresponding to residue types will be ignored' )
parser.add_argument( "-num_connections", type=int, default=48, help='Number of neighbors each residue is connected to, default 64, higher number leads to better interface design but will cost more to run the model.' )
parser.add_argument( "-fix_hotspot_res", action="store_true", help='Whether to fix the sequence of residues labelled as HOTSPOT or not (default False)' )

args = parser.parse_args( sys.argv[1:] )
silent = args.__getattribute__("silent")

omit_AAs = [ letter for letter in args.omit_AAs.upper() if letter in list("ARNDCQEGHILKMFPSTWYVX") ]

xml = "/home/nrbennet/protocols/xml/relax/dlpredictor/FastRelax/predictor_nodesign.xml"
objs = protocols.rosetta_scripts.XmlObjects.create_from_file( xml )

# Load the movers we will need

FastRelax = objs.get_mover( 'FastRelax' )
remove_massive_clashes = objs.get_mover( 'remove_massive_clashes' )
pack_no_design = objs.get_mover( 'pack_no_design' )
softish_min = objs.get_mover( 'softish_min' )
hard_min = objs.get_mover( 'hard_min' )
load_from_lite = objs.get_mover( 'load_from_lite' )
to_fa_standard = protocols.simple_moves.SwitchResidueTypeSetMover( 'fa_standard' )

ddg_filter = objs.get_filter( 'ddg' )
if ( isinstance(ddg_filter, pyrosetta.rosetta.protocols.filters.StochasticFilter) ):
    ddg_filter = ddg_filter.subfilter()

silent_out = "out.silent"

sfxn = core.scoring.ScoreFunctionFactory.create_score_function("beta_nov16")
objs.get_mover

alpha_1 = list("ARNDCQEGHILKMFPSTWYV-")
states = len(alpha_1)
alpha_3 = ['ALA','ARG','ASN','ASP','CYS','GLN','GLU','GLY','HIS','ILE',
           'LEU','LYS','MET','PHE','PRO','SER','THR','TRP','TYR','VAL','GAP']

aa_1_N = {a:n for n,a in enumerate(alpha_1)}
aa_3_N = {a:n for n,a in enumerate(alpha_3)}
aa_N_1 = {n:a for n,a in enumerate(alpha_1)}
aa_1_3 = {a:b for a,b in zip(alpha_1,alpha_3)}
aa_3_1 = {b:a for a,b in zip(alpha_1,alpha_3)}

# I/O Functions

def add2silent( pose, tag, sfd_out ):
    struct = sfd_out.create_SilentStructOP()
    struct.fill_struct( pose, tag )
    sfd_out.add_structure( struct )
    sfd_out.write_silent_struct( struct, "out.silent" )

# End I/O Functions

def my_rstrip(string, strip):
    if (string.endswith(strip)):
        return string[:-len(strip)]
    return string

def thread_mpnn_seq( pose, binder_seq ):
    rsd_set = pose.residue_type_set_for_pose( core.chemical.FULL_ATOM_t )

    for resi, mut_to in enumerate( binder_seq ):
        resi += 1 # 1 indexing
        name3 = aa_1_3[ mut_to ]
        new_res = core.conformation.ResidueFactory.create_residue( rsd_set.name_map( name3 ) )
        pose.replace_residue( resi, new_res, True )
    
    return pose

def sequence_optimize( pdbfile, chains, model, fixed_positions_dict ):
    
    t0 = time.time()

    feature_dict = mpnn_util.generate_seqopt_features( pdbfile, chains )

    seq_per_struct = 1
    arg_dict = mpnn_util.set_default_args( seq_per_struct, omit_AAs=omit_AAs )
    arg_dict['temperature'] = args.temperature

    masked_chains = [ chains[0] ]
    visible_chains = [ chains[1] ]

    sequences = mpnn_util.generate_sequences( model, device, feature_dict, arg_dict, masked_chains, visible_chains, fixed_positions_dict=fixed_positions_dict )
    
    print( f"MPNN generated {len(sequences)} sequences in {int( time.time() - t0 )} seconds" ) 

    return sequences

def get_final_dict(score_dict, string_dict):
    print(score_dict)
    final_dict = OrderedDict()
    keys_score = [] if score_dict is None else list(score_dict)
    keys_string = [] if string_dict is None else list(string_dict)

    all_keys = keys_score + keys_string

    argsort = sorted(range(len(all_keys)), key=lambda x: all_keys[x])

    for idx in argsort:
        key = all_keys[idx]

        if ( idx < len(keys_score) ):
            final_dict[key] = "%8.3f"%(score_dict[key])
        else:
            final_dict[key] = string_dict[key]

    return final_dict

def add2scorefile(tag, scorefilename, score_dict=None):
  
    write_header = not os.path.isfile(scorefilename)
    with open(scorefilename, "a") as f:
        add_to_score_file_open(tag, f, write_header, score_dict)

def add_to_score_file_open(tag, f, write_header=False, score_dict=None, string_dict=None):
    final_dict = get_final_dict( score_dict, string_dict )
    if ( write_header ):
        f.write("SCORE:     %s description\n"%(" ".join(final_dict.keys())))
    scores_string = " ".join(final_dict.values())
    f.write("SCORE:     %s        %s\n"%(scores_string, tag))

def generate_mut_string(seq):
    return 'MUT:' + '_'.join( [ f"{idx+1}.{aa_1_3[aa1]}" for idx, aa1 in enumerate(seq) ] )

def swap_mut_string(tag, mut_string, og_struct):
    outlines = []
    for line in og_struct:
        line = line.strip()
        if not 'PDBinfo-LABEL:' in line:
            # Swap out tags on all lines except for the remark line
            splits = line.split()
            if len(splits) == 0:
                outlines.append( '' )
                continue
            outline = splits[:-1]
            outline.append(tag)
            outlines.append( ' '.join(outline) )
            continue

        splits = line.split(' ')
        outsplits = []
        mut_found = False

        for split in splits:
            if not split.startswith('MUT:'):
                outsplits.append(split)
                continue
            mut_found = True
            outsplits.append(mut_string)

        if not mut_found: outsplits.append(mut_string)

        outlines.append( ' '.join(outsplits) )

    return '\n'.join(outlines)

def add2silent( pose, tag, sfd_out ):
    struct = sfd_out.create_SilentStructOP()
    struct.fill_struct( pose, tag )
    sfd_out.add_structure( struct )
    sfd_out.write_silent_struct( struct, "out.silent" )

def add2lite( seq, tag, og_struct ):
    
    mut_string = generate_mut_string(seq)
    new_struct = swap_mut_string(tag, mut_string, og_struct)
    with open( silent_out, 'a' ) as f: f.write( f"{new_struct}\n" ) 

def delete_file_list( filelist ):
    for f in filelist: os.remove( f )

def pdbs2poses( pdbs ):
    return [ ( pose_from_pdb( pdb ), my_rstrip( pdb, ".pdb" ) ) for pdb in pdbs ]

def get_chains( pose ):
    lengths = [ p.size() for p in pose.split_by_chain() ]
    endA = pose.split_by_chain()[1].size()
    endB = endA + pose.split_by_chain()[2].size()

    chains = [ pose.pdb_info().chain( i ) for i in [ endA, endB ] ]

    return chains

def get_hotspots(pose):
    hotspots = []
    pdb_info = pose.pdb_info()
    endA = pose.split_by_chain()[1].size()
    for i in range(1,endA+1):
        if pdb_info.res_haslabel(i,"HOTSPOT"):
            hotspots.append(i)
    return hotspots

def relax_pose( pose ):
    #remove_massive_clashes.apply( pose )
    FastRelax.apply( pose )

    return pose

def dlpredictor( pose ):

    remove_massive_clashes.apply( pose )
    pack_no_design.apply( pose )
    softish_min.apply( pose )
    hard_min.apply( pose )

    ddg = ddg_filter.compute( pose )
    
    return ddg

def dl_design( pose, tag, og_struct, mpnn_model, sfd_out ):

    tot_t0 = time.time()
    design_counter = 0

    prefix = f"{tag}_dldesign"
    pdbfile = f"tmp.pdb"

    if args.fix_hotspot_res:
        hotspots = get_hotspots( pose )
    else:
        hotspots = []
    
    fixed_positions_dict = None
    if len(hotspots)>0:
        fixed_positions_dict = {}
        fixed_positions_dict[my_rstrip(pdbfile,'.pdb')] = {"A":hotspots,"B":[]}
        print("Found residues with HOTSPOT label, fixing the following residues: ", fixed_positions_dict[my_rstrip(pdbfile,'.pdb')])

    for cycle in range(args.relax_cycles):
        pose.dump_pdb( pdbfile )
        chains = get_chains( pose )

        seqs_scores = sequence_optimize( pdbfile, chains, mpnn_model, fixed_positions_dict )
        os.remove( pdbfile )

        seq, mpnn_score = seqs_scores[0] # We know there is only one entry
        pose = thread_mpnn_seq( pose, seq )

        if args.output_intermediates:
            tag = f"{prefix}_0_cycle{cycle}"
            add2silent( pose, tag, sfd_out )
            design_counter += 1

        pose = relax_pose(pose)

    # Do the final sequence assignment
    pose.dump_pdb( pdbfile )
    chains = get_chains( pose )

    seqs_scores = sequence_optimize( pdbfile, chains, mpnn_model, fixed_positions_dict )
    os.remove( pdbfile )

    seq, mpnn_score = seqs_scores[0] # We know there is only one entry
    pose = thread_mpnn_seq( pose, seq )

    tag = f"{prefix}_0_cycle{args.relax_cycles}"
    if args.do_predictor:
        pose = relax_pose(pose)
        ddg = ddg_filter.compute(pose)
        add2scorefile( tag, 'out.sc', {'ddg':ddg, 'time':time.time()-tot_t0} )

    add2silent( pose, tag, sfd_out )

    design_counter += 1
    # Done with final sequence assignment

    return design_counter

def main( pdb, silent_structure, mpnn_model, sfd_in, sfd_out ):

    t0 = time.time()
    print( "Attempting pose: %s"%pdb )
    
    # Load pose from PDBLite
    pose = Pose()
    sfd_in.get_structure( pdb ).fill_pose( pose )
    if lite:
        load_from_lite.apply( pose )
        to_fa_standard.apply( pose )

    good_designs = dl_design( pose, pdb, silent_structure, mpnn_model, sfd_out )

    seconds = int(time.time() - t0)

    print( f"protocols.jd2.JobDistributor: {pdb} reported success. 1 designs generated in {seconds} seconds" )

def detect_lite(silentfile, silent_index):
    struct = silent_tools.get_silent_structure( silentfile, silent_index, silent_index['tags'][0] )
    for line in struct:
        if 'SCAFFOLD_PDB' in line: return True
    return False

# Checkpointing Functions

def record_checkpoint( pdb, checkpoint_filename ):
    with open( checkpoint_filename, 'a' ) as f:
        f.write( pdb )
        f.write( '\n' )

def determine_finished_structs( checkpoint_filename ):
    done_set = set()
    if not os.path.isfile( checkpoint_filename ): return done_set

    with open( checkpoint_filename, 'r' ) as f:
        for line in f:
            done_set.add( line.strip() )

    return done_set

# End Checkpointing Functions

#################################
# Begin Main Loop
#################################

silent_index = silent_tools.get_silent_index(silent)
lite = detect_lite(silent, silent_index) # using as a global variable

if not os.path.isfile(silent_out):
    with open(silent_out, 'w') as f: f.write(silent_tools.silent_header(silent_index))

sfd_out = core.io.silent.SilentFileData("out.silent", False, False, "binary", core.io.silent.SilentFileOptions())

sfd_in = rosetta.core.io.silent.SilentFileData(rosetta.core.io.silent.SilentFileOptions())
sfd_in.read_file(silent)

sf_open = open(silent)

checkpoint_filename = "check.point"
debug = True

if torch.cuda.is_available():
    print('Found GPU will run MPNN on GPU')
    device = "cuda:0"
else:
    print('No GPU found, running MPNN on CPU')
    device = "cpu"

finished_structs = determine_finished_structs( checkpoint_filename )
mpnn_model = mpnn_util.init_seq_optimize_model(device, hidden_dim=128, num_layers=3, backbone_noise=args.augment_eps, num_connections=args.num_connections, checkpoint_path=args.checkpoint_path)

for pdb in silent_index['tags']:

    if pdb in finished_structs: continue

    silent_structure = silent_tools.get_silent_structure_file_open( sf_open, silent_index, pdb )

    if debug: main( pdb, silent_structure, mpnn_model, sfd_in, sfd_out )

    else: # When not in debug mode the script will continue to run even when some poses fail
        t0 = time.time()

        try: main( pdb, silent_structure, mpnn_model, sfd_in, sfd_out )

        except KeyboardInterrupt: sys.exit( "Script killed by Control+C, exiting" )

        except:
            seconds = int(time.time() - t0)
            print( "protocols.jd2.JobDistributor: %s failed in %i seconds with error: %s"%( pdb, seconds, sys.exc_info()[0] ) )

    # We are done with one pdb, record that we finished
    record_checkpoint( pdb, checkpoint_filename )
```
