## Supplementary Information - Computational protocols and scripts for "De novo design of high-affinity miniprotein binders targeting *Francisella tularensis* virulence factor": polyXXX.py

```
#!/usr/bin/env python

from pyrosetta import *
from pyrosetta.rosetta import *

from argparse import ArgumentParser
import sys

def parse_arguments( argv ):
    argv_tmp = sys.argv
    sys.argv = argv
    description = "Mutate the whole protein to XXX !!!"
    parser = ArgumentParser( description = description )
    parser.add_argument('-pdbs', nargs="+", help='Input pdb file name')
    parser.add_argument('-residue', default='ALA', type=str, help='Target residue to mutate to, default to ALA')
    parser.add_argument('-gpc', default=False, action='store_true', help='Also change the Gly, Cys and Pro positions..')
    args = parser.parse_args()
    sys.argv = argv_tmp
    return args


def main( argv ):
    args = parse_arguments( argv )
    init()

    for pdb in args.pdbs:

        print(pdb)
        pose = pose_from_file( pdb )
        rts = rosetta.core.chemical.ChemicalManager.get_instance().residue_type_set("fa_standard")
        rtype = rts.name_map( args.residue )
        res = rosetta.core.conformation.ResidueFactory.create_residue( rtype )

        for ir in range(1, pose.size() + 1):
            name = pose.residue(ir).name3()
            if not args.gpc and ( name in ['GLY', 'PRO', 'CYS']):
                continue
            pose.replace_residue( ir, res, True )

        out = pdb.rstrip("gz").rstrip(".").rstrip("pdb").rstrip(".") + '_poly' + args.residue + '.pdb'
        pose.dump_pdb( out )


if __name__ == '__main__':
    main( sys.argv )
```
