## Supplementary Information - Computational protocols and scripts for "De novo design of high-affinity miniprotein binders targeting *Francisella tularensis* virulence factor": README.txt

############## SUPPLEMENTARY FILE DESCRIPTIONS ##############

rifgen.flag               -- Flag file for generating RIFs against selected target residues
auto_relax_beta_v3.1.sh   -- Script for automated PDB preparation and crystallographic refinement with Rosetta & Phenix
rifgen_residues.list      -- List of Flpp3 residues used for RIF generation
polyXXX.py                -- Script to standardize scaffold sequences by mutating all residues to VAL
predictor.flags           -- Example flag file for filtering RIFdock outputs prior to design
predictor_v11.xml         -- RosettaScript XML protocol for evaluating RIFdock poses prior to design
pilot.flags               -- Example flag file for pilot run selection and structural profiling
tutorial_design_v5.xml    -- RosettaScript XML protocol for pilot-stage design, refinement, and ML-driven filtering
dl_interface_design_R2.py -- Script for MPNN-guided protein interface design with structural refinement
predictor_nodesign.xml    -- RosettaScript XML protocol for post-design structural refinement and interface scoring
split_commands.py         -- Utility script for splitting large job command files for parallel SLURM array submission. It splits a shell command list into array-compatible job scripts. You may reuse or adapt this for any SLURM-based cluster.


############## DESIGN PIPELINE EXECUTION GUIDE ##############

# Make following directories:

PROJECT/rifdock_stage
PROJECT/fastdesign_stage1
PROJECT/grafting_stage
PROJECT/fastdesign_stage2
PROJECT/notebooks
PROJECT/to_order

# Run Rosetta and Phenix-based automated structure preparation and relaxation on input PDBs:
./auto_relax_beta_v3.1.sh

# Save your scaffold paths to a file named scaffolds.list

# Save your RIFgen residues to a file named rifgen_residues.list -one residue number per line, e.g. see example file-

# Edit rifgen.flag file. Set them as follows:

-rifgen:target – the path to your target pdb. This might just be target.pdb.
-rifgen:target_res – the path to your rifgen_residues.list file.
-rifgen:outdir – pick whatever name you want, probably rifgen_MYTARGET
-rifgen:outfile – any name works here, just make sure it ends with .rif.gz

# Generate RIFs using selected target residues. It is critical that you save the log:

OMP_NUM_THREADS=16 [YOUR_PATH_TO_LATEST_RIFGEN]/rifgen @rifgen.flag> rifgen.log 2>&1

# Make sure that your target chain is B
# Save your PatchDock interface residues to a file named patchdock_residues.list
# Format: one residue number per line, followed by the chain ID (e.g., 45 B)

# Convert the scaffolds to poly-valine:

[YOUR_PATH]/polyXXX.py -pdbs *.pdb -residue VAL

# Example on how to run it (supposing that your scaffold pdb paths are in a list called scaffolds.list):

mkdir poly_val_runs
mkdir poly_val_commands
cd poly_val_commands
split -l 1000 ../scaffolds.list
for j in x*; do echo "(cd $(readlink -f ../poly_val_runs); mkdir $j; cd $j; [YOUR_PATH]/polyXXX.py -pdbs \$(cat $(readlink -f $j)) -residue VAL)"; done> commands.list
[YOUR_PATH]/split_commands.py commands.list . 1 1
num_commands_min_1=$(( $(ls commands | wc -l) - 1))
sbatch -p short -a 0-$num_commands_min_1 array.submit

# Put your poly-valine scaffold paths into a list file called scaffolds_polyval.list.

# Setup your PatchDock jobs as described in: Cao, L.; Coventry, B.; et.al Design of Protein-Binding Proteins from the Target Structure Alone. Nature 2022, 605 (7910), 551–560.

# Setup your RifDock commands. You need to split the scaffolds into groups and pair them with their patchdock file. For these commands to work, it's critical that all of your patchdock outputs are in the folder called: PROJECT/rifdock_stage/patchdock_xforms.

RIFDOCK_FLAGS=rifdock.flag
PATCHDOCK_XFORMS=patchdock_xforms
SCAFFOLDS=scaffolds.list
RIFDOCK=[YOUR_PATH_TO_LATEST_RIFDOCK]/rif_dock_test
rifdock_flags=$(readlink -f $RIFDOCK_FLAGS)
patchdock_xforms=$(readlink -f $PATCHDOCK_XFORMS)
scaffolds=$(readlink -f $SCAFFOLDS)
mkdir rifdock_logs
mkdir rifdock_commands
cd rifdock_commands
mkdir splits
cd splits
split -l 40 $scaffolds
cd ..
mkdir patch_files
for j in splits/x*; do cat $j | sed -E 's%.*/%%g' | sed -E 's%.gz$%%g' | sed -E 's%.pdb$%%g' | sed  "s%^%$patchdock_xforms/%g" | sed 's%$%_polyVAL.out%g'> patch_files/$(basename $j); done
for j in splits/x*; do a=$(basename $j); echo "cd $(readlink -f ..);  $RIFDOCK @$rifdock_flags -scaffolds \$(cat rifdock_commands/splits/$a) -seeding_pos \$(cat rifdock_commands/patch_files/$a)> rifdock_logs/${a}.log 2>&1"; done> commands.list
cd ..

# Setup RIFdock jobs

[YOUR_PATH]/split_commands.py rifdock_commands/commands.list rifdock_commands 1   THE_AMOUNT_OF_RAM_YOU_NEED 15
num_commands_min_1=$(( $(ls rifdock_commands/commands | wc -l) - 1))
sbatch -p medium -a 0-$num_commands_min_1 rifdock_commands/array.submit

# Use the following command to get a list of all your rifdock outputs:

cat rifdock_out/*.silent> rifdock_output.silent

# You will need silenttools going forward: https://github.com/bcov77/silent_tools

# Designing RIFdock outputs:

RIFDOCK_OUTPUT=/<FULL_PATH_TO_YOUR_RIFDOCK_OUTPUT>/rifdock_out.silent
PREDICTOR_XML=predictor_v11.xml
PREDICTOR_FLAGS=predictor.flags
rifdock_output=$(readlink -f $RIFDOCK_OUTPUT)
predictor_xml=$(readlink -f $PREDICTOR_XML)
predictor_flags=$(readlink -f $PREDICTOR_FLAGS)
mkdir predictor_runs
mkdir -p predictor_commands/splits
cd predictor_commands/splits
silentsplitshuf $rifdock_output 200
for j in x*; do echo "cd $(readlink -f ../../predictor_runs); mkdir $j; cd $j;
[YOUR_PATH_TO_ROSETTA]/rosetta_scripts @$predictor_flags -parser:protocol $predictor_xml -in:file:silent $(readlink -f $j)> /dev/null 2>&1"; done>   ../commands.list
cd ../../

# Then submit the jobs with the following command:

[YOUR_PATH]/split_commands.py predictor_commands/commands.list predictor_commands 1 2
num_commands_min_1=$(( $(ls predictor_commands/commands | wc -l) - 1))
sbatch -p short -a 0-$num_commands_min_1 predictor_commands/array.submit

# Running the pilot jobs:

RIFDOCK_OUTPUT=../rifdock_stage/rifdock_out.silent
silentls $RIFDOCK_OUTPUT | shuf | head -n2000 > rifdock_for_pilot.list

RIFDOCK_OUTPUT=/<FULL_PATH_TO_YOUR_RIFDOCK_OUTPUT>/rifdock_out.silent
PILOT_XML=tutorial_design_v5.xml
PILOT_FLAGS=pilot.flags
rifdock_output=$(readlink -f $RIFDOCK_OUTPUT)
pilot_xml=$(readlink -f $PILOT_XML)
pilot_flags=$(readlink -f $PILOT_FLAGS)
mkdir pilot_runs
mkdir -p pilot_commands/splits
cd pilot_commands/splits
cat ../../rifdock_for_pilot.list | silentslice $rifdock_output> tmp.silent
silentsplitshuf tmp.silent 5
rm tmp.silent
for j in x*; do echo "cd $(readlink -f ../../pilot_runs); mkdir $j; cd $j;  
[YOUR_PATH_TO_ROSETTA]/rosetta_scripts @$pilot_flags -parser:protocol $pilot_xml -in:file:silent $(readlink -f $j)> /dev/null 2>&1"; done> ../commands.list
cd ../../

# Then submit the jobs with the following command:

[YOUR_PATH]/split_commands.py pilot_commands/commands.list pilot_commands 1 6
num_commands_min_1=$(( $(ls pilot_commands/commands | wc -l) - 1))
sbatch -p medium -a 0-$num_commands_min_1 pilot_commands/array.submit

# Collect predictor results by:

[YOUR_PATH]/fast_merge_scorefiles predictor_runs

# Collect pilot results by:

cat pilot_runs/*/out.silent> tmp.silent
silentls tmp.silent | silentrename tmp.silent> pilot_runs.silent
rm tmp.silent tmp.silent.idx
silentscorefile pilot_runs.silent

# Filter your designs based on your preference

# Setup your MPNN jobs (example to run one MPNN job):

cd [YOUR_PATH]/ppi_MPNN_FR_runs; mkdir xaa; cd xaa; [YOUR_PATH_TO_MPNN_ENV]/mpnn_binder_design.sif [YOUR_PATH]/dl_interface_design_R2.py -silent [YOUR_PATH]/ppi_MPNN_FR_commands/splits/xaa.silent -temperature 0.0001 -augment_eps 0 -relax_cycles 1 -checkpoint_path [YOUR_PATH_TO_PROTEINMPNN]/vanilla_model_weights/v_48_020.pt > log.log 2>&1

NOTE: v_48_020.pt can be obtained from the official ProteinMPNN GitHub repository: https://github.com/dauparas/ProteinMPNN

# Collect the results by:

cat ppi_MPNN_FR_runs/*/out.silent > collected.silent
