## Supplementary Information - Computational protocols and scripts for "De novo design of high-affinity miniprotein binders targeting *Francisella tularensis* virulence factor": split_commands.py

```
#!/usr/bin/env python

import os, sys, subprocess
import stat

group_size = int(sys.argv[3])
ramgb = int(sys.argv[4])
cpus = 1
if len(sys.argv) >= 6:
    cpus = int(sys.argv[5])


def cmd(command, wait=True):
    # print ""
    # print command
    the_command = subprocess.Popen(
        command, shell=True, stdout=subprocess.PIPE, stderr=subprocess.PIPE
    )
    if not wait:
        return
    the_stuff = the_command.communicate()
    return (the_stuff[0] + the_stuff[1]).decode("ascii")


folder = cmd("readlink -f %s" % sys.argv[2]).strip()
if not os.path.exists(folder):
    print("%s doesn't exist!!!" % folder)
    sys.exit(1)


to_open = sys.argv[1]

f = open(to_open)
lines = f.readlines()
f.close()

sets = []
this_set = []
for line in lines:
    line = line.strip()
    if len(line) == 0:
        continue
    this_set.append(line)
    if len(this_set) == group_size:
        sets.append(this_set)
        this_set = []

if len(this_set) > 0:
    sets.append(this_set)

if not os.path.exists("%s/logs" % folder):
    os.mkdir("%s/logs" % folder)

if not os.path.exists("%s/commands" % folder):
    os.mkdir("%s/commands" % folder)


count = 0
for this_set in sets:
    name = "%s/commands/command%i.sh" % (folder, count)
    f = open(name, "w")
    line = line.strip()
    f.write("#!/bin/bash\n")
    f.write("#SBATCH --mem=%ig\n" % ramgb)
    f.write("#SBATCH -N 1\n")
    f.write("#SBATCH -n 1\n")
    f.write("#SBATCH -c %i\n" % cpus)
    f.write("#SBATCH -o %s/logs/log%i.loglog\n" % (folder, count))
    f.write("\n".join(this_set))
    f.write("\n")
    f.close()
    count += 1

    st = os.stat(name)
    os.chmod(name, st.st_mode | stat.S_IXUSR | stat.S_IXGRP | stat.S_IXOTH)


with open("%s/array.submit" % folder, "w") as f:
    f.write("#!/bin/bash\n")
    f.write("#SBATCH --mem=%ig\n" % ramgb)
    f.write("#SBATCH -N 1\n")
    f.write("#SBATCH -n 1\n")
    f.write("#SBATCH -c %i\n" % cpus)
    f.write("#SBATCH -o %s/logs/log%%a.loglog\n" % (folder))
    f.write("\n")
    f.write("%s/commands/command${SLURM_ARRAY_TASK_ID}.sh\n" % (folder))
    f.close()
```
